# Microbially-catalyzed conjugation of GABA and tyramine to bile acids

**DOI:** 10.1101/2023.09.25.559407

**Authors:** Michael W. Mullowney, Aretha Fiebig, Matthew K. Schnizlein, Mary McMillin, Amber R. Rose, Jason Koval, David Rubin, Sushila Dalal, Mitchell L. Sogin, Eugene B. Chang, Ashley M. Sidebottom, Sean Crosson

## Abstract

Bile acids (BAs) are cholesterol-derived molecules that aid in digestion and nutrient absorption, regulate host metabolic processes, and influence physiology of the gut microbiota. Both the host and its microbiome contribute to enzymatic modifications that shape the chemical diversity of BAs in the gut. Several bacterial species have been reported to conjugate standard amino acids to BAs, but it was not known if bacteria conjugate BAs to other amine classes. Here, we show that *Bacteroides fragilis* strain P207, isolated from a bacterial bloom in the J-pouch of a patient with ulcerative colitis (UC) pouchitis, conjugates standard amino acids and the neuroactive amines γ-aminobutyric acid (GABA) and tyramine to deoxycholic acid. We extended this analysis to other human gut isolates and identified species that are competent to conjugate GABA and tyramine to primary and secondary BAs, and further identified diverse BA-GABA and BA-tyramine amides in human stool. A longitudinal metabolomic analysis of J-pouch contents of the patient from whom *B. fragilis* P207 was isolated revealed highly reduced levels of secondary bile acids and a shifting BA amide profile before, during, and after onset of pouchitis, including temporal changes in several BA-GABA amides. Treatment of pouchitis with ciprofloxacin was associated with a marked reduction of nearly all BA amides in the J-pouch. Our study expands the known repertoire of conjugated bile acids produced by bacteria to include BA conjugates to GABA and tyramine and demonstrates that these molecules are present in the human gut.

**Importance:** Bile acids (BAs) are modified in multiple ways by host enzymes and the microbiota to produce a chemically diverse set of molecules that assist in the digestive process and impact many physiological functions. This study reports the discovery of bacteria isolated from the gut of human patients that conjugate the neuroactive amines, GABA and tyramine, to BAs and demonstrates that BA-GABA and BA-tyramine amides are present in the human gut. GABA and tyramine are common metabolic products of the gut microbiota and potent neuroactive molecules, and their conjugation to BAs may influence receptor-mediated regulatory mechanisms of humans and their gut microbes.

## Introduction

Primary bile acids (PBAs) are produced from cholesterol in the liver through a multi-step enzymatic process (1). In the hepatocyte, PBAs are enzymatically conjugated to the amine group of glycine or taurine to form an amide bond (2, 3) and are eventually secreted into bile and released into the intestinal lumen. Once in the gut, members of the microbiota deconjugate glycine or taurine from PBAs and chemically modify the sterol core, yielding secondary bile acids (SBAs) of various chemical forms (4, 5). This chemically diverse ensemble of bile acids (BAs) in the human gut are eventually resorbed in the intestine and returned to the liver through enterohepatic circulation where they can be re-conjugated or repaired in the hepatocyte (6). Together, PBAs and SBAs in their various chemical forms play important roles in digestion and absorption of dietary fats, and regulate a range of host (1) and microbial (7) metabolic processes.

Recent studies have shown that the majority of standard protein-encoding amino acids can be conjugated to both PBAs and SBAs by bacteria that reside in the gut including species of the genera *Bacteroides, Lactobacillus, Bifidobacterium, Enterocloster, Ruminococcus*, and *Clostridium* (8-10). This discovery has expanded the known chemical repertoire of BAs present in humans, though the impact of these microbially-conjugated BAs on host and microbial physiology is not known. Certainly, the potent antibacterial properties of BAs have been recognized for over a century (11, 12), and it is long established that conjugated forms of BAs are less inhibitory to growth of intestinal microflora than unconjugated free acids *in vitro* (13-15). Gut bacteria may also encode protein receptors that cue specific regulatory responses to BAs including virulence gene expression (16, 17) and spore germination (18). The VtrA-VtrB-VtrC regulatory system of *Vibrio parahaemolyticus* is a notable example of the specificity of microbial responses to BAs. Though both conjugated and unconjugated bile acids bind the VtrA-VtrC receptor complex at the same site and with similar affinities (19), only specific BA species cue the transcription of the type III secretion regulator, *vtrB* (16, 17, 19). Beyond the effect of BAs on the microbiota, the impact of these molecules on mammalian physiology is well established and dependent on the chemical form of the molecule (20). For example, Takeda G protein-coupled receptor 5 (TGR5) and farnesoid X receptor (FXR) can function as bile acid receptors, and signaling through these receptors is influenced by the chemical identity of the amino acid that is conjugated to the sterol core (10, 21).

In ulcerative colitis (UC) patients who have undergone ileal pouch anal anastomosis (IPAA), a dysbiosis-induced deficiency of SBAs including DCA and lithocholic acid (LCA) is proposed to cue an inflammatory state that can lead to pouchitis (22). It is not known if microbial bile acid conjugation reactions directly modulate DCA and LCA levels in these patients, but genera with the capacity to conjugate amino acids to BAs (9), including *Bacteroides* spp., often dominate the inflamed ileoanal pouch (or J-pouch) (23). We have previously isolated *Bacteroides fragilis* strains that constitute over 50% of the bacterial population of the J-pouch before the emergence of inflammation (23). We aimed to test the ability of a predominant *B. fragilis* strain (P207) to conjugate amino acids and other amines present in the gut to primary and secondary bile acids. We further aimed to define the temporal bile acid profile of the J-pouch of the human patient from whom this particular *B. fragilis* strain was isolated.

Here, we report that *B. fragilis* strain P207 conjugates glycine (Gly), alanine (Ala), phenylalanine (Phe), γ-aminobutyric acid (GABA), and tyramine to DCA *in vitro*; conjugation to CA was limited to glycine. *B. fragilis* P207 deconjugates glycodeoxycholate (GDCA) to produce DCA, and can subsequently produce Ala-, Phe-, GABA-, and tyra-mine-DCA conjugates from the deconjugated bile acid. Thus, *B. fragilis* P207 produces a chemically diverse pool of conjugated bile acids *in vitro*, including the novel GABA-DCA and tyramine-DCA products. A time-series metabolomic analysis of stool from pouchitis patient 207 before and after onset of inflammation revealed severely diminished secondary bile acids across all time points and the presence of multiple bile acid-amine conjugates, the levels of which were strongly reduced following antibiotic treatment. Among the BA amides detected in pouch samples, or in samples from healthy human donors, were diverse GABA and tyramine conjugates. GABA- and tyra-mine-BA amide synthesis was not limited to *B. fragilis* P207, as we identified other classes of gut anaerobes that conjugate these neuroactive amines to PBAs and SBAs. Our results expand the known set of microbially-catalyzed bile conjugation reactions and have identified novel bile acid conjugates to GABA and tyramine in the human gut. Both GABA (24) and tyramine (25) are common products of microbial metabolism and are potent neuromodulators. Flux of these amines in the gut due to microbe-catalyzed conjugation to bile acids may impact host physiology.

## Results

### Chemical transformation of primary and secondary bile acids by *B. fragilis* P207

*B. fragilis* P207 was incubated in supplemented Brain Heart Infusion (BHIS) broth with and without deoxycholic acid (DCA) (0.01% w/v), and samples were analyzed by ultra-high-performance liquid chromatography tandem high resolution mass spectrometry (UHPLC-MS^2^). Fragmentation-based networking of the mass spectrometry data identified derivatives of DCA from a set of candidate conjugates presented in Table S1. This metabolomic approach provided evidence for five amide-linked conjugates of DCA, which were only present in the spent media of cultures that contained both *B. fragilis* P207 and DCA. Fragmentation patterns consistent with DCA-alanine, DCA-glycine, DCA-phenylalanine, DCA-tyramine, and DCA-γ-aminobutyric acid (DCA-GABA) were identified (Figure 1). We aimed to validate the synthesis of bile acid-amine conjugates by *B. fragilis* P207 and to investigate if this strain could link exogenous amines by adding isotopically labeled variants of these compounds to the culture medium. Specifically, we supplemented broth containing *B. fragilis* P207 and deoxycholic acid (0.01% w/v) with 1 mM of either isotopically labeled 4-aminobutyric acid (^13^C_4_, 97-99%), L-alanine (^13^C_3_, 99%), glycine (^13^C_2_, 99%; ^15^N, 98%+), L-phenylalanine (D_8_, 98%), or tyramine:HCl (1,1,2,2-D_4_, 98%). Analyses of spent media from each of these conditions revealed the expected intact mass (MS^1^) and fragments (MS^2^) for all five DCA conjugates (<0.48 ppm error for all MS^1^ ions and <6 ppm error for all MS^2^ ions representing predicted structures), as well as the expected mass shifts when media was supplemented with the isotopically labelled precursors (Figure 2; Figures S1–S5). Given that bile acid conjugation to GABA and tyramine have not been previously reported, we further validated the identity of these compounds by comparing their LC retention times and MS data to synthetic standards. These experiments further confirmed the identity of these bile acid amides produced by *B. fragilis* P207 as DCA-GABA and DCA-tyramine (Figure 3 and Figures S6-S7). Mass spectrometry evidence for production of these conjugates is fully presented in Supplemental Results.

**Figure 1.**
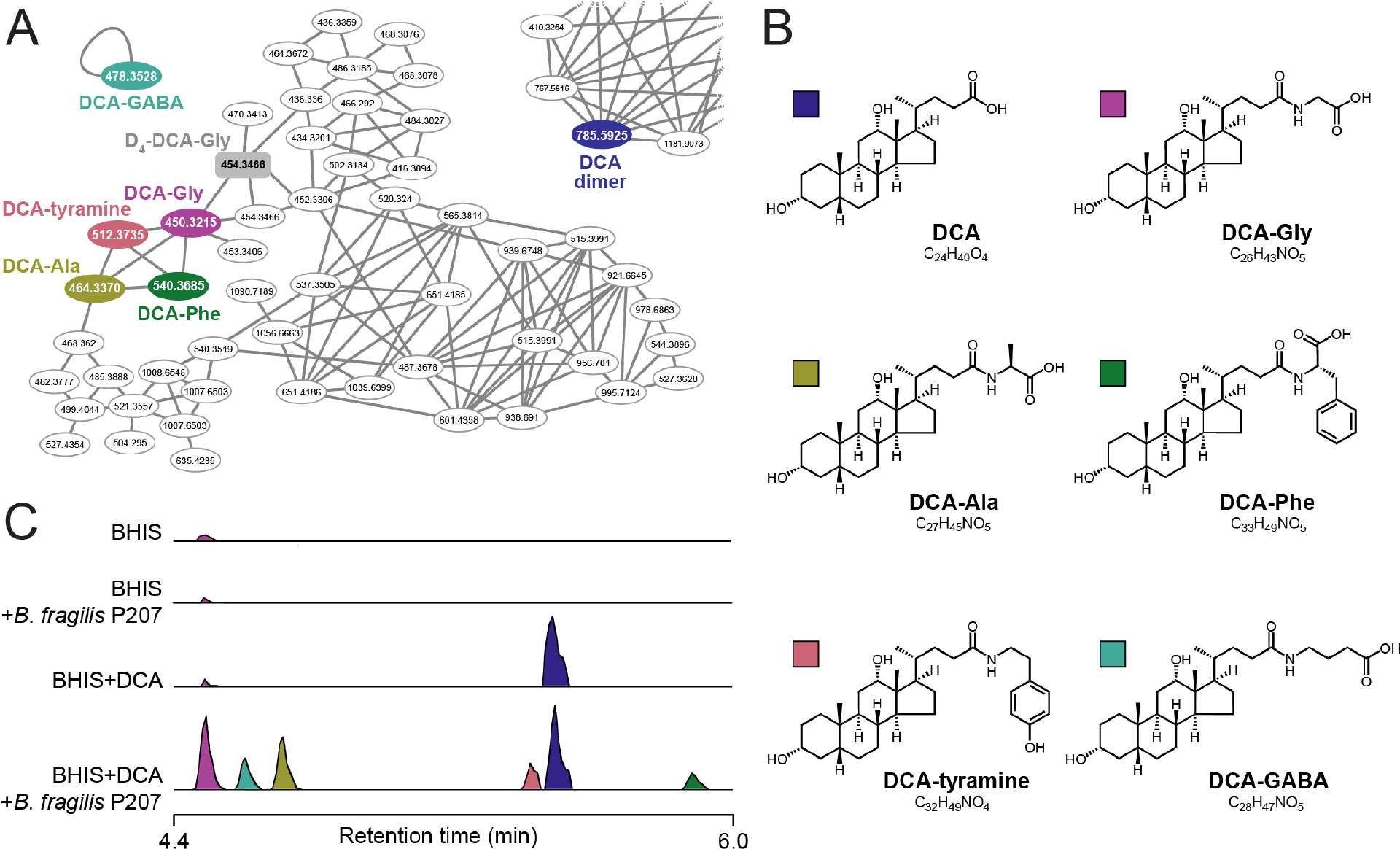
Identification of bile acid conjugates produced by pouchitis patient isolate, *B. fragilis* P207 by UHPLC-MS^2^ (**A)** Molecular family containing nodes for deoxycholic acid (DCA)-amine conjugates. Colored nodes represent features only observed in *B. fragilis* P207 cultures with 0.01% (w/v) deoxycholic acid (DCA) present. The gray-shaded rectangular node is the D_4_-DCA-Gly internal standard. **(B)** Structures of the five DCA-amine conjugates with color key for panels A, B, and C. (**C)** Stacked selected ion chromatograms for the five DCA-amino acid conjugates detected in BHIS media (BHIS), *B. fragilis* strain P207 culture extract (BHIS+*B. fragilis* 207), BHIS media with 0.01% DCA (BHIS+DCA), and *B. fragilis* strain P207 culture extract with DCA (BHIS+*B. fragilis* P207+DCA).

**Figure 2.**
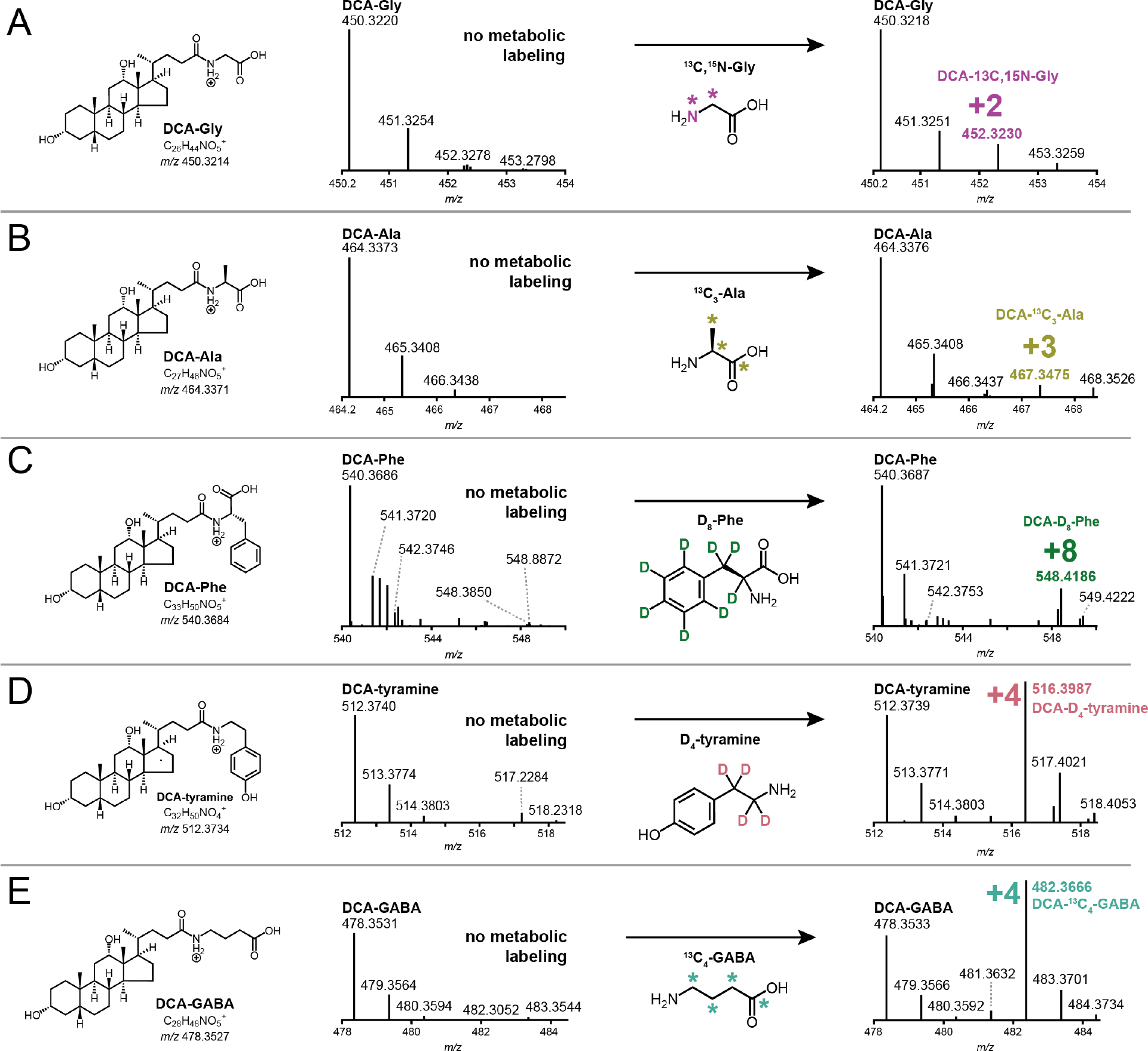
Amine feeding experiments to test bile acid conjugation by *B. fragilis* strain P207. Mass spectrometry analysis was performed to assess production of bile acid conjugates with five different amines: glycine (A), alanine (B), phenylalanine (C), tyramine (D), and GABA (E). *B. fragilis* P207 was fed isotopically labeled versions of these amines. Observed mass to charge (m/z) shifts, displayed in each panel, are consistent with the expected molecular weights for the respective bile acid-heavy amino acid/amine conjugates presented in Figure 1.

**Figure 3.**
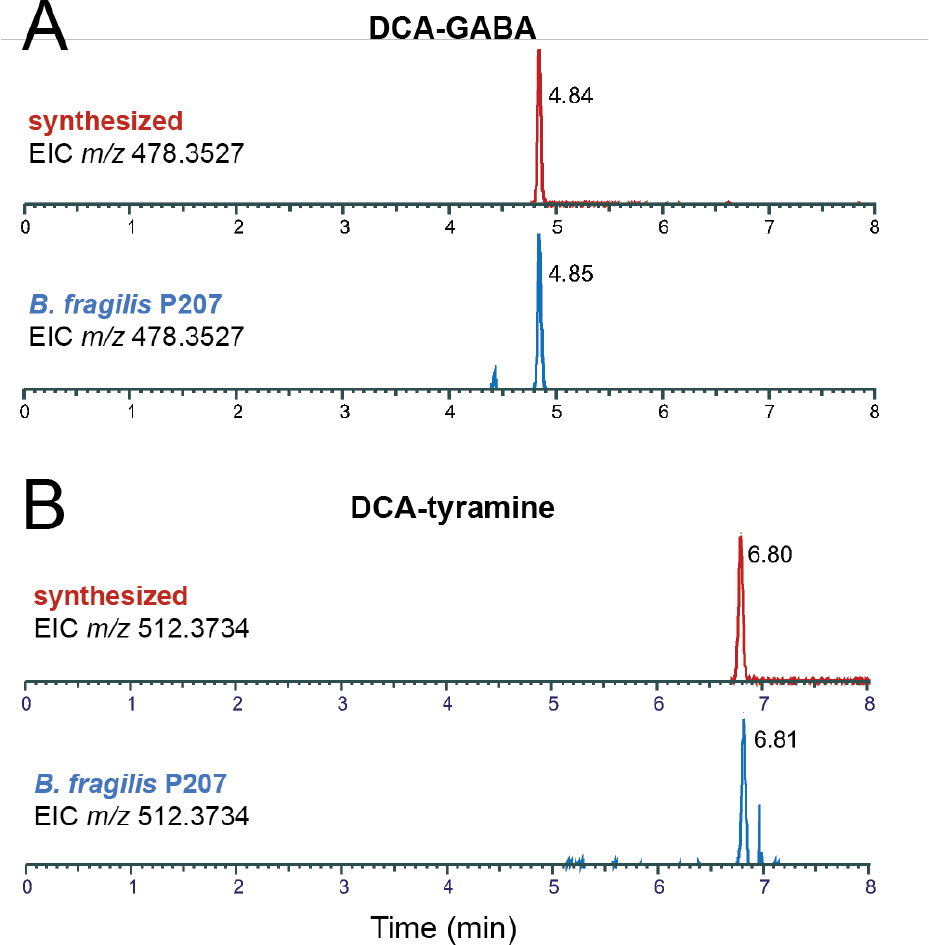
Extracted ion chromatograms (EICs) of deoxycholate conjugates produced by *B. fragilis* P207 *in vitro* compared to synthetic standards. (A) EIC of chemically synthesized DCA-GABA (m/z 478.3527) (top; retention time=4.84 minutes) and the EIC of DCA-GABA identified in *B. fragilis* P207 *in vitro* cultures (bottom; retention time=4.85 minutes). (B) EIC of chemically synthesized DCA-tyramine (m/z 512.3734) (top; retention time=6.80 minutes) and the EIC of DCA-tyramine identified in *B. fragilis* P207 *in vitro* cultures (bottom; retention time=6.81 minutes). After identifying bile acid conjugates as key metabolites (Figure 1), we refined the chromatography method to effectively separate the complete array of possible conjugate isomers, considering both deoxycholic acid (DCA) and cholic acid (CA) isomers as additional potential substrates. This accounts for the differences in retention times from Figure 1.

To assess chemical specificity of bile acid conjugation by *B. fragilis* P207, we further tested for production of cholic acid (CA) amides. Mass spectrometry analyses revealed trace levels of DCA in the BHIS growth media and traces of DCA and GCA in the starting CA reagent; GCA levels were enhanced by incubation of CA (0.01% w/v) with *B. fragilis* P207 (Figure 4A). Simultaneous incubation of CA and DCA (0.01% w/v total) with strain P207 showed similar production of the same five DCA conjugates described above. Thus, the addition of CA did not apparently impact P207-dependent production of the five DCA conjugates over this timescale (Figure 4B and Table S2). We conclude that strain P207 prefers DCA as a conjugation substrate over CA in BHIS medium.

**Figure 4.**
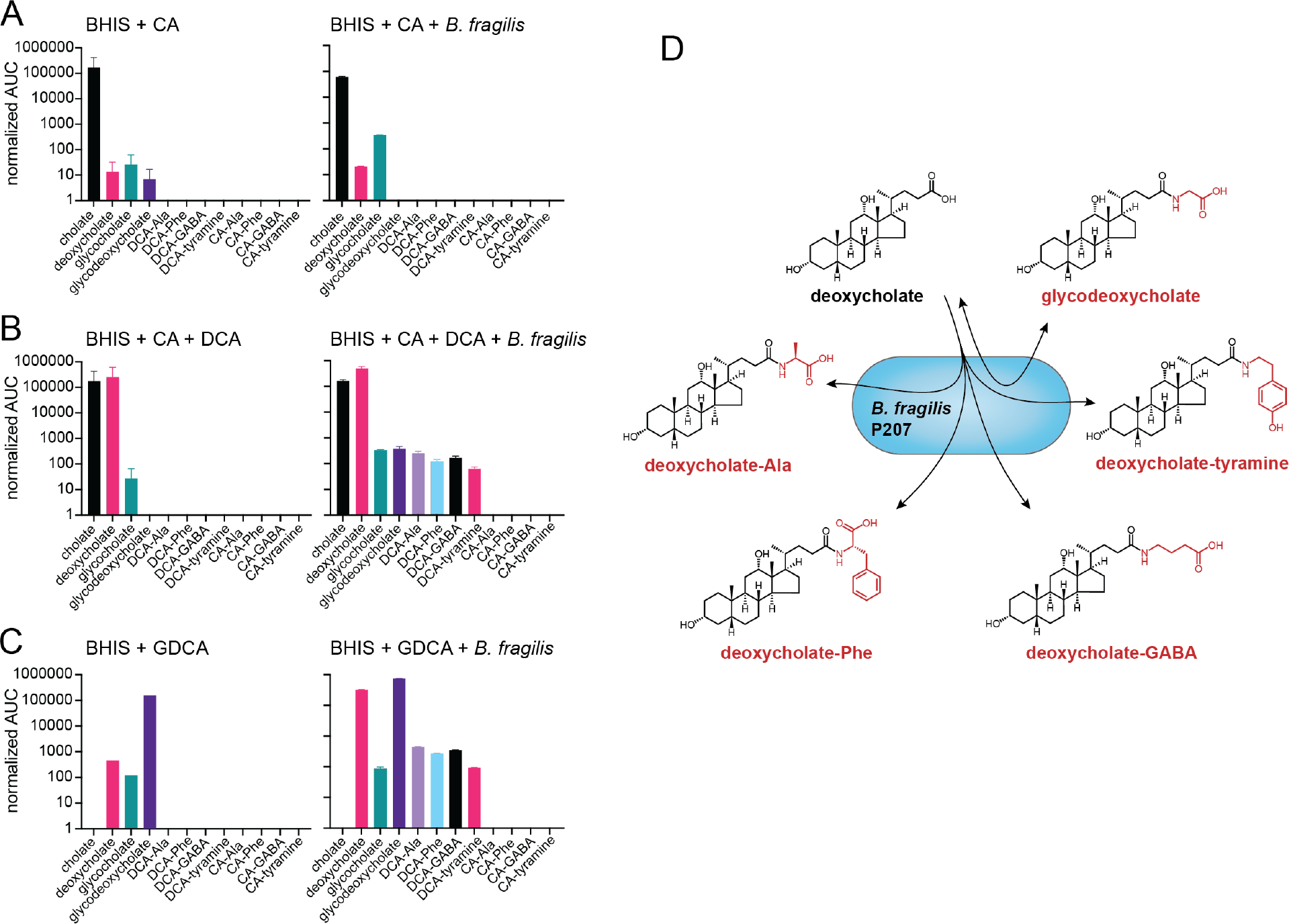
LC-MS/MS measurements of amino acid/amine conjugation to the bile acids cholic acid (CA) and deoxycholic acid (DCA), and evidence for deconjugation/re-conjugation of glycodeoxycholic acid (GDCA) by *B. fragilis* P207. Bar graphs represent the mean area under the curve (AUC) of LC-MS/MS chromatographic peaks corresponding to unconjugated or conjugated bile acids. (A) AUC of bile acid conjugate peaks (labeled below each bar) when *B. fragilis* P207 was cultivated in BHIS broth the presence of 0.01% (w/v) CA. (B) AUC of bile acid conjugate peaks when *B. fragilis* P207 was cultivated in BHIS broth the presence of CA and DCA (0.01% (w/v) total). (C) AUC of bile acid and bile acid conjugate peaks from *B. fragilis* P207 culture extract cultivated in BHIS broth the presence of 0.01% (w/v) GDCA. Cholate and deoxycholate are known contaminants of the GDCA reagent. Data represent the mean ± SD of two independent cultures. The absence of a bar indicates that a peak corresponding to that chemical species was not detected. (D) Cartoon representing the bile acid conjugate production profile of the human gut isolate, *B. fragilis* P207.

Lucas and colleagues have demonstrated that modification of CA into the secondary bile acid, 7-oxoDCA, is robust and widespread across several species of *Bacteroides*, while production of DCA from CA was limited to *Bacteroides vulgatus* (9). We did not observe increased DCA levels when *B. fragilis* P207 was incubated with CA, nor did we observe production of any DCA conjugates from CA indicating that *B. fragilis* P207 does not catalyze production of DCA from CA under these conditions. The relative abundances of bile acid amide conjugates produced across replicate experiments are presented in Figure S8.

### P207 catalyzed deconjugation and reconjugation of amines to DCA

Given that some bacteria can deconjugate glycine and taurine from secreted bile acids, we tested whether *B. fragilis* P207 is able to produce the DCA conjugates described above using glycodeoxycholate (i.e. GDCA) as an initial substrate. *B. fragilis* P207 was incubated in BHIS broth with and without GDCA (0.01% w/v), and culture samples were analyzed by UHPLC-MS^2^. We observed trace contaminants DCA and glycocholic acid (GCA) in the growth medium and bile acid reagent, but incubation of GDCA with strain P207 resulted in decreased GDCA and increased DCA levels providing evidence that *B. fragilis* P207 has GDCA deconjugation activity (Figure 4C). We further observed production of the DCA conjugates described above (GABA, tyramine, alanine, phenylalanine) from GDCA when *B. fragilis* was present in the medium (Figure 4C). We conclude that *B. fragilis* P207 can produce all the DCA conjugates described in the section above when either DCA or GDCA is present as an initial substrate.

### GABA production by *B. fragilis* P207 is induced by deoxycholate

The observation of bile acid conjugation to a distinct set of amines *in vitro* raised the question about the levels of these amines in the culture medium. To measure the relative levels of specific amine-containing compounds in *B. fragilis* P207 cultures before and after exposure to DCA, we used gas chromatography-mass spectrometry (GC/MS) to analyze the same culture samples in which the DCA conjugates were identified. Among the five amines conjugated to DCA, alanine and phenylalanine were most abundant in the medium, followed by glycine at levels 15- to 40-times lower. Tyramine and GABA were present at levels approximately 10 times lower than glycine (Figure 5). Relative amine abundances did not, therefore, directly correlate with the levels of their respective DCA conjugates produced *in vitro* (Figure S8).

**Figure 5:**
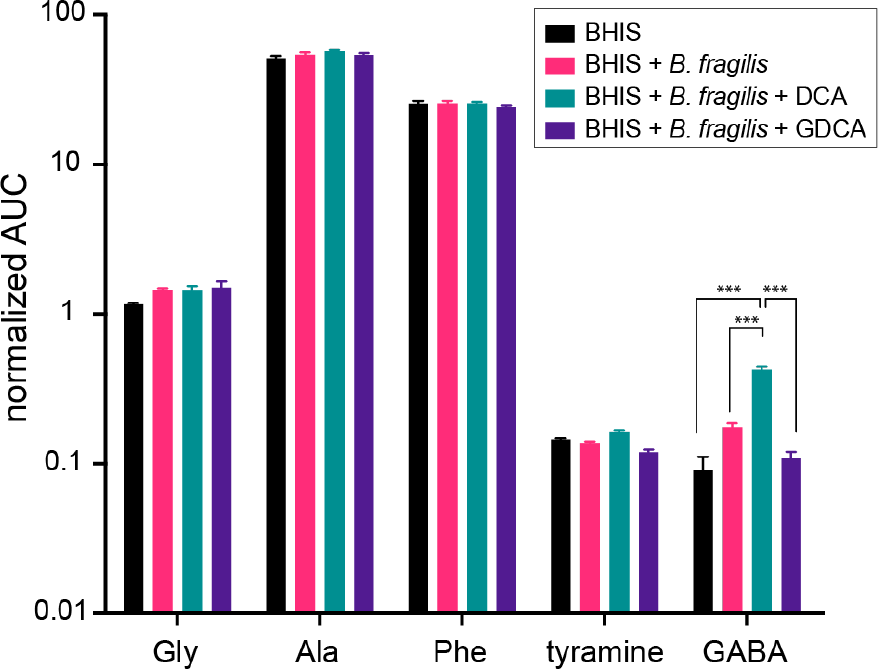
GC/MS-based detection of tyramine, glycine, alanine, phenylalanine, and GABA in *B. fragilis* P207 cultures. Cultures were grown in BHIS medium, both in the absence and presence of bile acids, DCA, and GDCA (0.01%, w/v). The displayed values, derived from the area under the curve (AUC) of detected peaks, represent the relative concentrations of these amine compounds across the different cultures. Each bar represents the mean of three independent experiments, with error bars indicating the standard deviations. Differences between conditions were assessed for statistical significance using ANOVA with Bonferroni correction. A threshold of p < 0.001 was considered to indicate statistical significance, ***.

Notably, when DCA was added to the culture medium, a significant increase in GABA was observed; glycodeoxycholic acid (GDCA) addition did not have this effect on microbial GABA production. The relative concentrations of other conjugated amines (i.e. alanine, phenylalanine, tyramine) was unchanged across all treatment conditions (Figure 5 and Figure S9). *Bacteroides* species in the human gut are known to produce GABA from either glutamate or glutamine, particularly under pH stress conditions (26). Given the increase in GABA following DCA treatment, we expanded our GC/MS analysis to include glutamine, glutamate, as well as other naturally occurring amino acids. DCA treatment did not impact glutamine levels, but it did lead to a modest reduction in glutamate concentrations (q < 0.01; Figure S9), which is consistent with a model whereby DCA stimulates *B. fragilis* P207 to convert glutamate into GABA. However, DCA was recently reported to strongly induce expression of glutamate dehydrogenase (gene locus *PTOS_003163*) in *B. fragilis* P207 (27), suggesting that the metabolite flux involving glutamate is likely modulated in multiple ways by DCA. Cultivation of *B. fragilis* P207 did not greatly affect the levels of amino acids in BHIS overall, though glycine, GABA, glutamine, histidine, methionine, proline and tyrosine increased significantly (q < 0.01) after *B. fragilis* was cultivated in the medium. Asparagine and aspartate showed the largest changes in the presence of *B. fragilis*: the concentration of these two amino acids decreased by approximate factors of 50 and 10, respectively (q < 0.001), suggesting they serve as primary nutritional substrates for *B. fragilis* P207 in BHIS medium (Figure S9).

### Defining the bile acid conjugate profiles of a pouchitis patient and healthy donors

To test if the bile acid conjugates produced by *B. fragilis* P207 *in vitro* were present in the gut of the human patient from whom the strain was isolated, we prepared extracts of stool samples collected from patient 207 at timepoints before and after onset of pouchitis, and during antimicrobial treatment (23). Metabolomic data from this patient were compared to UHPLC-MS^2^ data from 21 healthy donor stool samples. Patient 207 and healthy donor stool contained a complex mixture of bile acids (Figure 6 and Table S2). Unlike healthy donors, which contained expected high levels of DCA, patient 207 lacked DCA across all time points (Figure S10 and Table S2). This result is congruent with a reported deficiency in secondary bile acids in UC pouches (28). The primary bile acids chenodeoxycholate (CDCA) and CA were abundant across all time points in patient 207, at levels that were ≈60 times higher on average than observed in the 21 healthy donor samples (p < 0.001) (Figure S11 and Table S2). Amine conjugates to CA and CDCA, including putative GABA conjugates, were identified in patient 207 stool (Figures 6-7; Figures S12-S13 and Table S2). We detected several amines conjugated to a bile acid core with an *m/z* corresponding to DCA or its isomers (Figure 6 and Figure S8). Considering the lack of DCA and abundance of CDCA in patient 207, we predicted that these species were CDCA derivatives. Indeed, a product with an elution profile and MS^2^ fragmentation pattern matching a CDCA-GABA synthetic standard was identified in patient 207 stool (Figure 7 and Figure S12). We further identified products matching CA-GABA and CA-tyramine synthetic standards in patient 207 stool and in healthy human donor stool, respectively (Figure 7 and Figures S13-S14). The results provide strong evidence that bile acids conjugated to GABA and tyramine are present in the human gut.

**Figure 6.**
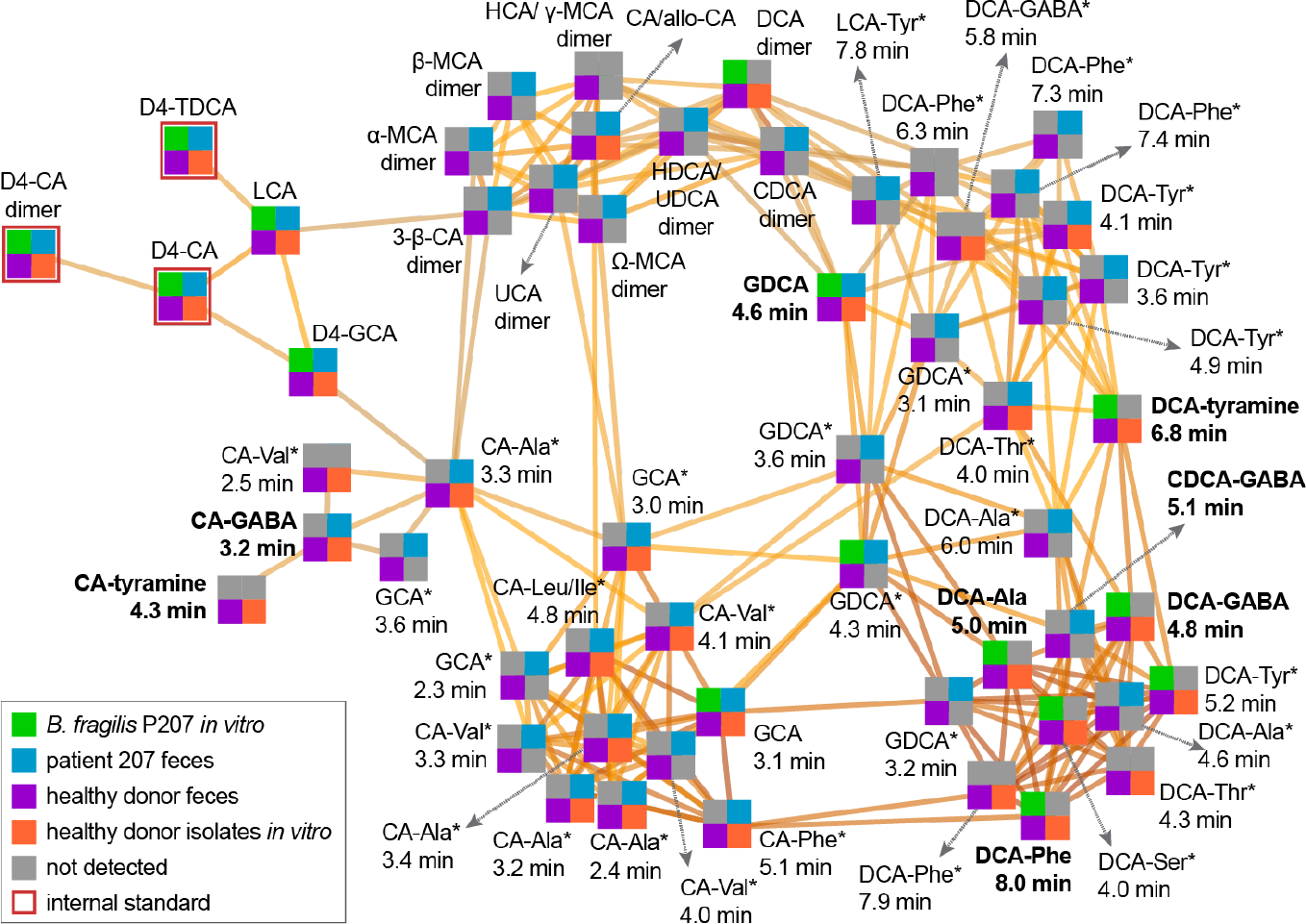
Molecular network illustrating the diversity and occurrence of bile acid-amine conjugates related to the validated DCA-Phe, DCA-Ala, DCA-Gly, DCA-tyramine, DCA-GABA, CDCA-GABA, CA-GABA, and CA-tyramine products. Each node represents a high-resolution *m/z* value at a specific retention time. Node quadrant color indicates the sample type that the metabolite was detected in—gray color indicates that the metabolite was not detected in the sample type specified by that quadrant. Edges connect nodes that are related above a score threshold of 25 in Compound Discoverer software suite (Thermo Scientific), with darker edges signifying greater relatedness. Bolded node labels indicate metabolites validated by isotope labeling (GDCA, DCA-Ala, DCA-Phe) and/or comparison to true synthetic standards (DCA-GABA, DCA-tyramine, CDCA-GABA, CA-GABA, CA-tyramine). Labels with an asterisk (*) indicate that the node represents a putative bile acid-amine isomer/epimer of the listed metabolite. These isomer assignments are based on comparison of *m/z* of the intact ion and MS^2^ fragmentation spectra and fragment *m/z* values to the spectra of the validated bile acid conjugates. All other metabolite nodes were confirmed by comparison to authenticated standards or labeled internal standards. Chenodeoxycholic acid (CDCA); deoxycholic acid (DCA); cholic acid (CA); glycocholic acid (GCA), lithocholic acid (LCA); taurodeoxycholic acid (TDCA); ursocholic acid (UCA); hyodeoxycholic acid (HDCA); ursodeoxycholic acid (UDCA); hyocholic acid (HCA); muricholic acid (MCA). D4 indicates deuterated standards.

**Figure 7.**
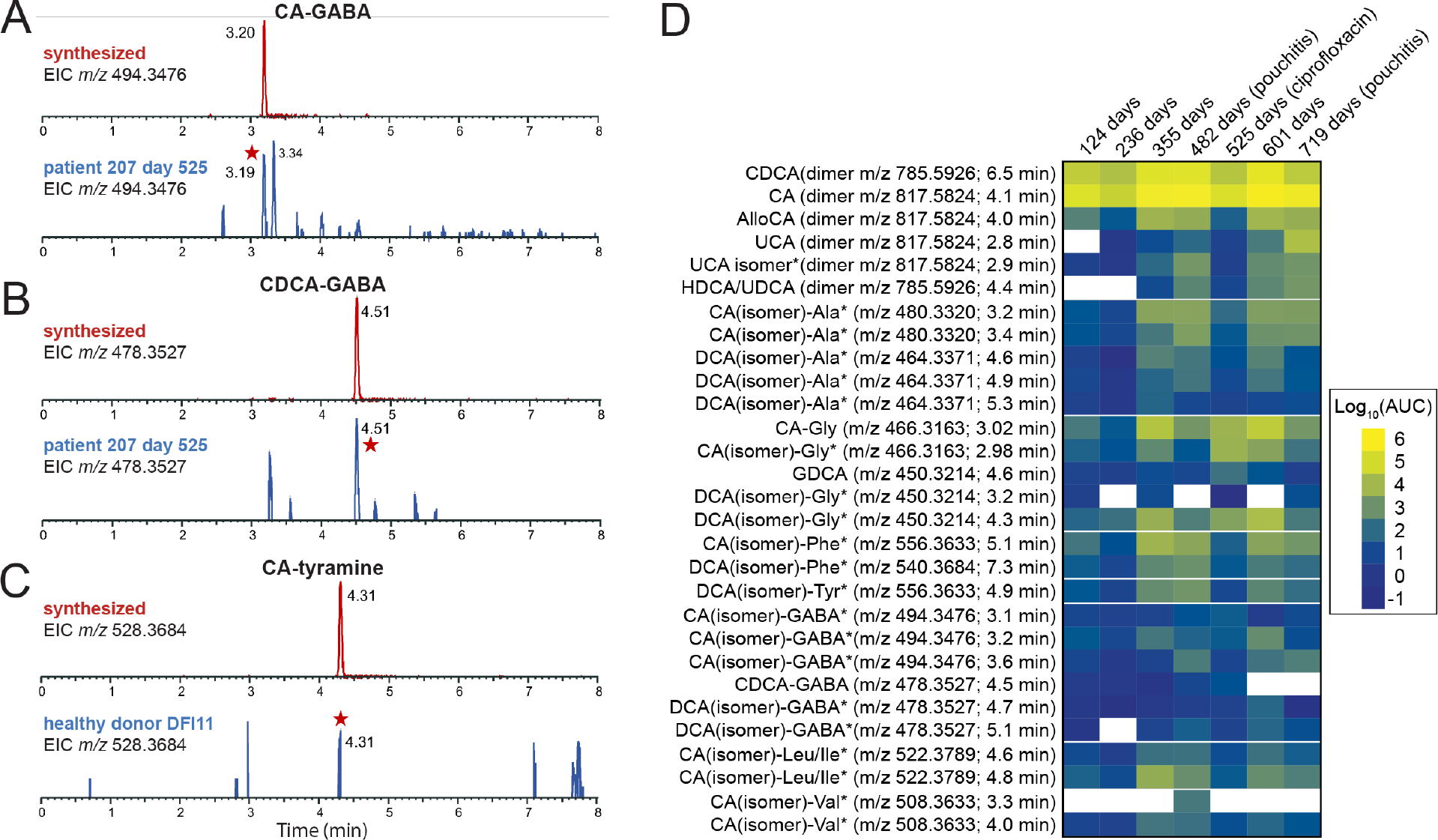
Extracted ion chromatograms (EICs) of synthetic standards of cholic acid (CA) or chenodeoxycholic acid (CDCA) conjugated to GABA or tyramine compared to stool sample extracts. (A) EIC of chemically synthesized CA-GABA (*m/z* 494.3476) (top; retention time=3.20 minutes) and the corresponding EIC from patient 207 stool at 525 days post functionalization of the ileal pouch (bottom). Peak matching synthetic CA-GABA is marked with a red star. (B) EIC of chemically synthesized CDCA-GABA (*m/z* 478.3527) (top; retention time=4.51 minutes) and the corresponding EIC from patient 207 at 525 days post functionalization of the ileal pouch (bottom). Peak matching CDCA-GABA is marked with a red star. (C) EIC of chemically synthesized CA-tyramine (*m/z* 528.3864) (top; retention time=4.31 minutes) and the corresponding EIC from healthy patient donor (DFI11) stool (bottom). Peak matching synthetic CA-tyramine is marked with a red star. (D) Heatmap illustrating the levels of unconjugated and conjugated bile acids in pouchitis patient 207 (measured by area under the curve or AUC) from days 124 to 719 after J-pouch functionalization. DCA (3α,12α-Dihydroxy-5β-cholan-24-oic acid) was absent at all time points in this patient and CDCA was abundant, so we expect the labeled DCA(isomer) amide conjugates have a CDCA core, though the isomeric/epimeric form is not defined in most cases. Likewise, we expect that many of the abundant CA(isomer) conjugates have a cholic acid core but the particular isomer/epimer of these products has not been defined.

*B. fragilis* strain P207 dominated the pouch ecosystem of patient 207 from 182 to 434 days after surgical functionalization of the ileal J-pouch (i.e. IPAA) (23). Normalized levels of bile acids, including the primary bile acids CA and CDCA, were lowest at 124 days and 236 days after IPAA. There was a marked increase in unconjugated CA and CDCA and bile acid conjugates at 355 days post-functionalization, and levels trended upward by 482 days when the patient was diagnosed with pouchitis and initiated a course of antibiotic therapy (ciprofloxacin) (Figure 7D and Figure S11). The contribution of *B. fragilis* strain P207 to the production of particular bile acid conjugates at these time points is not known, but the *in vitro* conjugation data presented above indicate that CA conjugates (other than glycocholate) are not produced by *B. fragilis* P207 (Figure 4). At a follow-up visit 525 days post-IPAA, ciprofloxacin treatment had resulted in a marked reduction of the pouch microbiome census and pouch inflammation was resolved (23). Bile acid analysis showed that levels of conjugated and unconjugated bile acids were sharply reduced at this time point except for select low abundance conjugates of CA and DCA isomers to glycine and GABA, which increased (Figure 7D, Figures S11 and S15 and Table S2). Though inflammation was resolved by ciprofloxacin treatment, inflammation returned by 719 days (23). In this period, the dominant species of the pouch shifted from *B. fragilis* P207 to *Bacteroides ovatus* (23). This was correlated with a change in the bile acid conjugate profile, with distinct CA and DCA isomer conjugates to GABA and glycine increasing (Table S2, Figures S11 and S15)

### Conjugation of GABA and tyramine to CA by human gut isolates

Metabolomic analysis of stool samples from pouchitis patient 207 and healthy human donors revealed a complex mixture of known and previously unreported bile acid conjugates, including CA conjugates to GABA and tyramine (Figures 6-7). To discover bacteria that can produce these novel CA conjugates, we inspected the genomes of a collection of human gut isolates for genes that encode predicted N-terminal nucleophilic cysteine hydrolase (Ntn) enzymes, which includes known choloylglycine hydrolases (Conserved Domain Database accession cd01902; (29)). Recent studies report that this class of enzymes can function to both deconjugate and conjugate bile acids (30, 31). *B. fragilis* P207 encodes an Ntn hydrolase (WP_005817456.1) and we predicted that other strains encoding these enzymes may conjugate GABA or tyramine to CA *in vitro*. We selected a phylogenetically diverse group of human patient isolates that were also predicted to encode an Ntn hydrolase, including *Mediterraneibacter gnavus* MSK15.77 (WP_004614568.1), *Bifidobacterium longum* DFI.2.45 (WP_007052221.1), and *Bacteroides ovatus* MSK22.29 (which encodes three Ntn paralogs: WP_217723859.1, WP_004308262.1, and WP_004323538.1) (Figure S15). We further identified a strain in our collection that does not encode a predicted Ntn bile salt hydrolase, *Lachnoclostridium scindens* SL.1.22. All three Ntn-encoding gut isolates produced putative GABA conjugates to CA, and *M. gnavus* and *B. longum* cultures contained a product that had an MS^2^ fragmentation pattern consistent with CA-tyramine (Figure 8 and Figure S16). *L. scindens* cultures did not contain CA conjugates.

**Figure 8.**
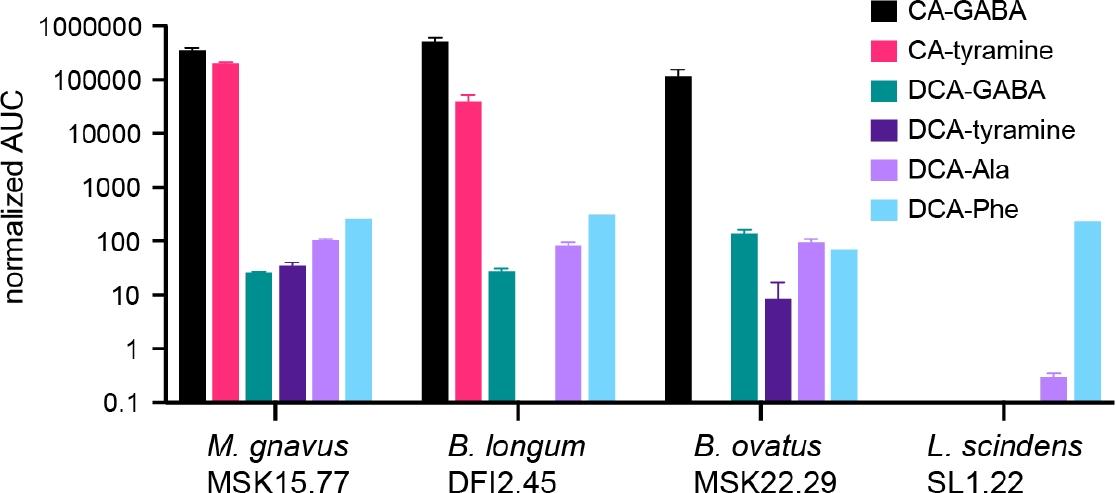
UHPLC-MS^2^ measurements of amino acid/amine conjugation to the bile acids cholic acid (CA) and deoxycholic acid (DCA) by *M. gnavus, B. longum, B. ovatus*, and *L. scindens* strains isolated from healthy human patients. The bar graphs in this figure represent the mean area under the curve (AUC) of LC-MS/MS peaks corresponding to unconjugated or conjugated CA and DCA (n=2, errors bars are standard deviation). Normalized AUC of bile acid conjugate peaks (colored according to key) when strains were cultivated in BHIS broth the presence of 0.01% (w/v) DCA or CA. The absence of a bar indicates that a peak corresponding to that chemical species was not detected.

We further investigated if these strains conjugated amines to DCA and detected conjugates of alanine, phenylalanine, and GABA to DCA in the culture extracts of *M. gnavus, B. longum*, and *B. ovatus*. Additionally, DCA-tyramine conjugates were identified in *M. gnavus* and *B. ovatus* cultures (Figure 8). Although the *L. scindens* SL.1.22 genome does not encode a predicted Ntn family/bile salt hydrolase (BSH) enzyme, we observed low levels of DCA-Phe and DCA-Ala conjugates in its culture extract when this strain was incubated with 0.01% (w/v) DCA. The specific gene(s) responsible for DCA conjugation in our *L. scindens* cultures remain unidentified.

## Discussion

### On the mechanism of microbial bile acid conjugation

*B. fragilis* P207 produces a suite of DCA amides *in vitro*, in which the bile acid carboxylate group is conjugated to the amine group of select amino acids and neuroactive amines. Biosynthesis of bile acid amides by gut microbes is a recent discovery (9, 10), and there is now genetic and biochemical evidence that bacterial bile salt hydrolase (BSH) enzymes can catalyze aminoacyl transfer to bile salts (30, 31). Thus, BSH proteins have significant N-acyl transferase activity in addition to their long-established function as hydrolases (32). While microbial conjugation of bile acids to the α amino group of amino acids has been described, our data provide evidence that an activity encoded by P207 enables conjugation of the primary amine groups of GABA and tyramine to DCA. DCA conjugate biosynthesis was evident when culture broth was supplemented with either unconjugated (DCA) or conjugated (GDCA) bile acids, which supports a model in which P207 first deconjugates GDCA and then generates DCA amides from the deconjugated product.

The *B. fragilis* P207 genome (GenBank accession CP114371) encodes a single predicted BSH (gene locus *PTOS_003312*) that shares 80% identity with BT2086, a protein that has demonstrated BSH activity in *B. thetaiotaomicron* (33). PTOS_003312 is 99% identical to the BSH of *B. fragilis* strain 638R (locus *BF638R_3310*), which promotes deconjugation of primary bile acids in the gut of germ-free mice (34). In light of recent *in vitro* biochemical data showing that purified *Clostridium perfrigens* (30) and *Bifidobacterium longum* (31) BSH enzymes produce bile acid amides from both unconjugated (CA) and conjugated (taurocholic acid; TCA) bile acids, it seems most likely that PTOS_003312 of *B. fragilis* P207 catalyzes the production of the five bile acid conjugates reported here, using either GDCA or DCA as substrates, though we cannot rule out the possibility that these products arise from the activity of multiple enzymes. P207 demonstrates a clear preference for bile acid amide production from the secondary bile acid, DCA, over primary bile acid CA, which differ only by a single hydroxyl group at C-7. Based on this result, we infer that the C-7 position of the sterol core is important for bile acid substrate interaction with the transforming enzyme(s) of strain P207.

The detection of DCA conjugation to the primary amine groups of GABA and tyramine raises questions about the mechanism of bile acid conjugation given the relative difference in nucleophilicity and steric accessibility of the primary amine groups of tyramine and GABA (pKa ≈ 10.5-11.0 in aqueous solvent) and the typical α amino group (pKa ≈ 9-9.5) of amino acids. Considering the diversity in primary structure of BSH enzymes and the selectivity of BSH enzymes for particular steroidal cores across *Bacteroides* spp. (33), it is likely that structural differences in the active site or other regions of the conjugating enzyme (35) will determine whether primary amine groups, such as those on GABA and tyramine, form amide bonds with bile acid(s). Evidence for microbially-catalyzed production of GABA and tyramine BA amides presented herein is congruent with a recent report that a diverse array of amines in the gut are conjugated to bile acids (36).

The benefit of bile acid conjugation to *B. fragilis* and other microbes studied here, if any, is not known. *B. fragilis* P207 can deconjugate GDCA and produce a variety of conjugated compounds from the deconjugated bile acid substrate (DCA) (Figure 4). Unconjugated primary and secondary bile acids are often more toxic *in vitro* than the amino acid-conjugated forms (13, 37, 38). Indeed, we have shown that *B. fragilis* P207 is more sensitive to CA and DCA than glycocholate and glycodeoxycholate (Figure S17). We further observed that the secondary bile acid, DCA, is more toxic to *B. fragilis* P207 than CA. The ability to conjugate amino acids effectively detoxifies DCA *in vitro* and may therefore influence the impact of bile salts on *B. fragilis* physiology in certain settings.

### Broader connections of bile acid conjugation to mammalian physiology

This study defines the temporal bile acid profile of the J-pouch of a pouchitis patient, before, during, and after inflammation including a period of antibiotic therapy, and draws comparisons of this profile to stool of healthy individuals. Healthy human donors presented expected high levels of DCA, while the pouchitis patient was severely DCA deficient. This aligns with previous observations of reduced secondary bile acids in UC pouches (28). As expected for a patient with diminished secondary bile acid production, the primary bile acids, CDCA and CA, were significantly elevated and unique amine conjugates to CA and CDCA, particularly novel GABA conjugates, were identified in this patient. *In vitro* analyses of *B. fragilis* P207 and of other human gut microbiome isolates has demonstrated that bile acid conjugates to GABA and tyramine observed in stool can be produced by multiple bacterial genera. Changes in the levels of bile acids leading up to pouchitis, coupled with the marked influence of ciprofloxacin treatment on the bile acid profile (Figures S9 and S12), underscore the complex interplay between gut microbiota, host factors, and therapeutic interventions on the bile acid profile of humans.

The neuroactive amines, GABA and tyramine, are potent regulators of mammalian physiology. Both molecules are produced by microbes that inhabit the gut microbiome, including Bacteroidetes (26). GABA has a well-recognized role in the regulation of gut physiology (39). GABA-producing *Bacteroides* increase steady-state levels of GABA in the intestines of mono-associated germ-free mice (40). We have shown that bile acid exposure enhances microbial GABA production *in vitro* (Figure 5) and provided evidence that exogenous GABA can be conjugated by B. fragilis P207 (Figure 2). The conjugation and subsequent uptake of bile acid-GABA amides into enterohepatic circulation may impact GABAergic signaling in the gut and possibly at other distal tissue sites. Tyramine is a product of tyrosine catabolism but is also present at high levels in a variety of foods. Steady-state tyramine levels in human tissue are typically low due to the activity of monoamine oxidase A, but can be modulated in patients treated with monoamine oxidase inhibitors (41). Trace amine associated receptors (TAARs), with which tyramine can interact, have been reported in enterochromaffin (EC) cells of the human gut (42), though the impact of dietary and microbiome-derived tyramine of TAAR signaling in the gut is not known. Tyramine-bile acid amides in human patients likely vary as a function of diet and other factors and vary geographically along the digestive tract depending on the amount of microbial tyramine production and the spectrum of bile acid conjugating activity conferred by the host microbiome at particular foci. Tyramine feeding data presented in Figure 2 suggest that dietary tyramine could impact BA-tyramine levels in the gut.

The impact of specific conjugated bile acids reported here on signaling through bile acid receptors is an interesting future area of investigation. Bile salt hydrolase (BSH) activity of non-enterotoxigenic *Bacteroides* spp. was recently reported to potentiate obesity-related colorectal cancer progression in a mouse model (43); this effect was attributed in part to enhanced TGR5 signaling as a result of increased levels of deconjugated DCA and lithocholic acid in the colon. Certainly, gut microbe BSH activity can lower the level of deconjugated bile acids in the gut. However, it is important to consider that BSH activity results in a spectrum of unconjugated and conjugated bile acids (30, 31), including compounds reported here. The effect of these compounds on signaling from bile acid receptors may shape host physiology, disease, and health.

## Materials and Methods

### Cultivation of bacteria in the presence of bile acids

*B. fragilis* P207 (NCBI locus NZ_CP114371) was cultivated in Brain-Heart infusion medium supplemented with hemin (BHIS) containing either 0.01% (w/v) deoxycholic acid (DCA), cholic acid (CA) or glycodeoxycholic acid (GDCA) that was inoculated from a saturated starter culture ( ≈1.0 OD_600_), back-diluted 1:10. Cultures were grown for 24 hours; 1 ml of culture was removed and flash frozen for subsequent metabolomic analysis. *Mediterraneibacter gnavus* MSK15.77 (NCBI accession NZ_JAAIRR010000000), *Bacteroides ovatus* MSK22.29 (NCBI accession NZ_JAHOCX010000000), *Bifidobacterium longum* DFI.2.45 (NCBI accession NZ_JA-JCNS010000000), and *Lachnoclostridium scindens* SL.1.22 (NCBI accession GCA_020555615.1) were also cultivated in BHIS. Briefly, starter cultures were inoculated from frozen glycerol stocks and grown anaerobically at 37°C overnight to ≈1.0 OD_600_. These cultures were diluted to ≈0.005 OD_600_ in 1 ml of plain BHIS or BHIS containing 0.01% (w/v) DCA or CA in a 96-well deep well plate. The plate was incubated at 37°C anaerobically for 20 hours and then frozen at -80°C for subsequent metabolomic analysis.

### Untargeted metabolomic approach to detect bile acid conjugates

Bacteria culture supernatants were lyophilized followed by 2X concentration in 100% methanol (containing internal standards). Samples were then centrifuged at −10 °C, 20,000 x *g* for 15 min to generate supernatants for subsequent metabolomic analysis. All ultra-high-performance-liquid chromatography-mass spectrometry (UHPLC-MS) analyses were performed using a Thermo Scientific Vanquish Flex UHPLC coupled with an IQ-X mass spectrometer (Thermo Fisher Scientific). Reversed-phase chromatography was performed on a Waters CORTECS T3 C18 RP-UHPLC column (100 × 2.1 mm inner diameter, 1.6 μm particle size, 120 Å pore size (1 Å = 0.1 nm)). Mobile phase A was 5% acetonitrile in water with 0.1% formic acid and mobile phase B was acetonitrile with 0.1% formic acid. For the initial analysis of *B. fragilis* P207 cultures (as displayed in Figure 1), the flow rate was 550 μL • min^-1^. The chromatographic method used was an isocratic 20% mobile phase B for 0.2 min, followed by a gradient of 20 to 97% mobile phase B for 8.7 min with a wash of 97% mobile phase B for 1.0 min. Analysis of *B. fragilis* P207 cultures for normalized relative quantification (as displayed in Figure S6) was performed at a flow rate of 480 μL • min^-1^. The chromatographic method used was an isocratic 100% mobile phase A for 0.2 min, followed by a gradient of 0 to 97% mobile phase B for 10.2 min with a wash of 97% mobile phase B for 1.0 min. Optimized chromatography was then developed for all other HUPLC-MS/MS experiments to better separate the numerous isomers of the various bile acid amine conjugates. The chromatographic method used a flow rate of 300 μL • min^-1^ with a gradient of 20 to 40% mobile phase B for 0.2 min, followed by a gradient of 40 to 70% mobile phase B for 11.8 min, a gradient of 70 to 97% mobile phase B for 0.1 min, and a wash of 97% mobile phase B for 1.1 min.

All samples were analyzed using positive ionization. Flow from the UHPLC was ionized with the heated electrospray ionization (HESI) source set to 3400 V, ion transfer tube temp set to 200 °C, vaporizer temperature set to 400 °C, and sheath, aux and sweep gases set to arbitrary values of 40, 5, and 1, respectively. Data for MS^1^ were acquired using a maximum inject time of 50 ms, a normalized AGC target of 25%, a 300–1700 m/z quadrupole scan range, and a resolution of 60,000. All tandem MS^2^ mass spectral data were acquired using a 1.5 m/z quadrupole isolation window, a maximum inject time of 22 ms, a normalized AGC target of 20%, and a resolution of 15,000. Each of the metabolite ions of interest were added to an inclusion list for MS^2^ fragmentation by a normalized higher-energy collisional dissociation energy of 30%. Data analysis was performed using FreeStyle software (version 1.8 SP2, Thermo Scientific), MZmine 2.53 (44) GNPS platform tools Feature-Based Molecular Networking (FBMN) release 28.2 and Mass Search Tool (MASST) release 29, (45-47), and GraphPad Prism version 9.4.1 for Windows (GraphPad Software, San Diego, California, USA, www.graphpad.com). Areas under the curve (AUC) were normalized to the average of D_4_-taurocholate and D_4_-taurodeoxycholate areas under the curve and multiplied by 1000:

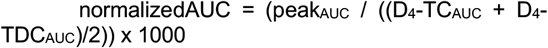

### Data processing and molecular networking

Data-dependent mass spectrometry data files were processed using MZmine 2.53 and the Feature-Based Molecular Networking function in the Global Natural Products Social Molecular Networking (GNPS) environment to identify and network spectral features while also matching generated data to publicly available library spectra. MZmine was used to detect MS^1^ and MS^2^ peaks, build deconvoluted extracted ion chromatograms, group isotopes, and match the resulting features across samples, accounting for retention time drift across injections. Features within each sample were then normalized by dividing peak area by the average peak area of the internal standards in that sample. Features that are found in solvent and method blank controls were then eliminated. The aligned feature lists were then exported for analysis using the FBMN tool within the GNPS platform.

FBMN was performed with precursor ion mass tolerance and fragment ion mass tolerance both set to 0.02 Da, minimum matched fragment ions set to six, networking cosine score set to > 0.7, library cosine score set to > 0.7, and minimum library shared peaks set to six. A visualization of the network was constructed in Cytoscape by drawing edges between scan nodes with a cosine similarity > 0.7. The network was manually analyzed to identify ions that occurred only in samples containing BHIS, a bacterial strain, and DCA or CA, with a particular focus on those that occurred within the same molecular families as our bile acid internal standards and other nodes that matched to library entries of known bile acids. Nodes in the molecular network were also checked against a list of predicted *m/z* values for hypothesized amine-containing conjugates presented in Table S1. Data files were also manually analyzed for MS^2^ scans containing characteristic DCA core fragments at *m/z* 215.1794 and *m/z* 339.2682 (observable in Figures S1-S5) to identify metabolite ions that may not have been clustered in the molecular network due to falling below networking thresholds.

### Isotopically labeled amine feeding experiments

To test for conjugation of isotopically labeled amines to DCA by *B. fragilis* P207, 1 ml cultures containing 1 mM (final concentration) of either 4-aminobutyric acid (^13^C4, 97-99%), L-alanine (^13^C3, 99%), glycine (2-^13^C, 99%; ^15^N, 98%+), L-phenylalanine (D_8_, 98%), or tyramine:HCl (1,1,2,2-D_4_, 98%) (Cambridge Isotope Laboratories, Inc.) was added to BHIS containing 0.1% (w/v) deoxycholic acid (DCA) (Fisher Scientific). Control samples contained either no isotopically labeled amines, no DCA, or BHIS media without any additional supplement. Culture supernatants were prepared for metabolomic analysis as described above.

### Synthesis of bile acid conjugate standards

Synthesis of bile acid–amine conjugates followed methods previously described (10). Cholic acid, deoxycholic acid, and chenodeoxycholic acid were dissolved to 0.125 mM in 5 mL anhydrous THF, each in two separate 20 mL scintillation vials containing stir bars and kept on an ice bath, for a total of six vials. Next, ethyl chloroformate (0.15 mmol, 7 µL) and triethylamine (0.15 mmol, 10.5 µL) were added to each reaction vessel and stirred for 1.5 h in the ice bath. Basic aqueous solutions of GABA (1.54 eq, 0.193 mM) and tyramine (1.54 eq, 0.193 mM) were prepared using NaOH (1.5 eq, 0.188 mM), with 5 mL of the GABA solution added to the first of each pair of bile acid solutions and 5 mL of the tyramine solution added to the second of each pair of bile acid solutions. The reaction mixtures were stirred for 2 h, centrifuged at −10 °C, 20,000 x g for 15 min to remove any remaining insoluble reaction components, and injected on the UHPLC-MS/MS using the optimized bile acid chromatography method.

### Metabolite analysis using GC-EI-MS with methoxyamine and TMS derivatization

Amine-containing metabolites were measured using GCMS with Electron Impact Ionization. Bacterial cultures were extracted using four volumes of 100% methanol. Following brief vortexing and centrifugation at 4°C, 20,000 x g for 15 min, 100 µL of extract supernatant was added to prelabeled mass spec autosampler vials (Microliter; 09-1200) and dried down completely under nitrogen stream at 30 L/min (top) 1 L/min (bottom) at 30°C (Biotage SPE Dry 96 Dual; 3579M). To dried samples, 50 µL of freshly prepared 20 mg/mL methoxyamine (Sigma; 226904) in pyridine (Sigma; 270970) was added and incubated in a thermomixer C (Eppendorf) for 90 min at 30°C and 1400 rpm. After samples are cooled to room temperature, 80 µL of derivatizing reagent (BSTFA + 1% TMCS; Sigma; B-023) and 70 µL of ethyl acetate (Sigma; 439169) were added and samples were incubated in a thermomixer at 70°C for 1 hour and 1400rpm. Samples were cooled to RT and 400 µL of Ethyl Acetate was added to dilute samples. Turbid samples were transferred to microcentrifuge tubes and centrifuged at 4°C, 20,000 x g for 15 min. Supernatants were then added to mass spec vials for GC-MS analysis. Samples were analyzed using a GC-MS (Agilent 7890A GC system, Agilent 5975C MS detector) operating in electron impact ionization mode, using a HP-5MSUI column (30 m x 0.25 mm, 0.25 µm; Agilent Technologies 19091S-433UI) and 1 µL injection. Oven ramp parameters: 1 min hold at 60°C, 16° C per min up to 300° C with a 7 min hold at 300°C. Inlet temperature was 280° C and transfer line was 300° C. Data analysis was performed using MassHunter Quantitative Analysis software (version B.10, Agilent Technologies) and confirmed by comparison to authentic standards. Normalized peak areas were calculated by dividing raw peak areas of targeted analytes by averaged raw peak areas of ^15^N,d_7_-L-proline and U-^13^C-palmitate internal standards.

### Bile acid analysis of human stool

Patient 207 feces was liquid and extraction was performed by vortexing and diluting 1:2 (200 µL into 400 µL) with 100% methanol containing internal standards, followed by bath sonication for 10 minutes. Non-diseased volunteer donor feces were solid and were extracted by adding 80% methanol to 100 mg/mL and stored at −80 °C for at least one hour in beadruptor tubes (Fisherbrand; 15-340-154). Donor samples were then homogenized at 4 °C on a Bead Mill 24 Homogenizer (Fisher; 15-340-163) set at 1.6 m/s with six thirty-second cycles, five seconds off per cycle. All samples were then centrifuged at −10 °C, 20,000 x *g* for 15 min to generate supernatants for subsequent metabolomic analysis. Areas under the curve for culture supernatants were normalized to the average of D_4_-taurocholate and D_4_-taurodeoxycholate areas under the curve and multiplied by 1000, while areas under the curve for fecal samples were normalized to the average of D_4_-taurocholate and D_4_-glycocholate areas under the curve and multiplied by 1000. All peak assignments in stool samples were verified by comparing the MS^2^ fragmentation spectra to MS^2^ fragmentation spectra established from the *in vitro* analyses described above.

### B. fragilis P207 bile sensitivity growth assays

*B. fragilis* strain P207 was cultivated in BHIS supplemented with increasing concentrations of DCA, CA, GDCA, or GCA. Concentrations ranged from 0.024 mM to 6.17 mM of each compound. Strain P207 was grown from a saturated starter culture (∼1.0 OD_600_) that was back-diluted 1:100. Cultures were then grown for 24 hours, at which point terminal culture density was measured at OD_600_.

### Mass spectrometry data availability

All mass spectrometry data files for this study have been submitted to the MassIVE data repository (https://massive.ucsd.edu/) in raw and open source formats under accession numbers MSV000093027, MSV000093028, MSV000093029, MSV000093030, MSV000093031, MSV000093032, MSV000093035, and MSV000093039. The GNPS Feature-Based Molecular Networking (FBMN) job used to aid in the identification of the deoxycholic acid amine conjugates from the initial untargeted screen of *B. fragilis* P207 cultures can be accessed on the GNPS platform (https://gnps.ucsd.edu/ProteoSAFe/status.jsp?task=2911fb25124d4a2784a72e720b57cfcc)

## Acknowledgements

We thank Doug Guzior and Rob Quinn for helpful discussion and references. Research reported in this publication was supported in part by the Duchossois Family Institute and the National Institutes of Health awards 5RC2DK122394 to E.C. and R35GM131762 to S.C.

## Supplemental Results and Figures

### Mass spectrometry data

#### Glycine conjugate

Broth from P207 cultures containing DCA and unlabeled glycine or isotopically labeled glycine (^13^C_2_, 99%; ^15^N, 98%+) revealed a DCA-Gly conjugate with an [M+H]^+^ of *m/z* 450.3220 and a corresponding labeled ion with an [M+H]^+^ of *m/z* 452.3230, respectively (Figure 2A). Analysis of tandem MS fragmentation indicated condensation of the amine of Gly with DCA at the terminal side-chain carboxylic acid (Figure S1). Key MS^2^ fragments from unlabeled P207 culture samples that indicate the position of the glycine on DCA include the doubly dehydroxylated DCA-Gly conjugate ion (*m/z* 414.3007), the *N*-(1-Oxo-4-penten-1-yl)glycine ion (*m/z* 158.0807), and the glycine aminium ion (*m/z* 76.0393). Ions with two Dalton increases corresponding to these were observed in MS^2^ data associated with an MS^1^ scan with a two Dalton increase in the glycine (^13^C_2_, 99%; ^15^N, 98%+) labeled sample, further confirming the structural assignment of glycine conjugation at the terminal side-chain carboxylic acid of DCA.

#### Alanine conjugate

Broth from strain P207 cultures containing DCA and unlabeled L-alanine or isotopically labeled L-alanine (^13^C_3_, 99%) revealed a DCA-Ala conjugate with an [M+H]^+^ of *m/z* 464.3373 and a corresponding labeled ion with an [M+H]^+^ of *m/z* 467.3475, respectively (Figure 2B). The tandem MS fragmentation profile indicated condensation of the amine of Ala with DCA at the terminal side-chain carboxylic acid (Figure S2). Key MS^2^ fragments from unlabeled P207 cultures that support the position of the alanine on DCA include the doubly dehydroxylated DCA-Ala conjugate ion (*m/z* 428.3165), the *N*-(1-Oxo-4-penten-1-yl)alanine ion (*m/z* 172.0963), and the alanine aminium ion (*m/z* 90.0544). Ions with three Dalton increases corresponding to the dehydroxylated DCA-Ala conjugate and the alanine aminium ion were observed in MS^2^ data associated with an MS^1^ scan with a three Dalton increase in the L-alanine (^13^C_3_, 99%) sample. These results further support the structural assignment of alanine conjugation at the terminal side-chain carboxylic acid of DCA, which was recently described using an alternative derivatization method (1).

#### Phenylalanine conjugate

Comparison of broth from P207 cultures containing DCA and unlabeled L-phenylalanine or isotopically labeled L-phenylalanine (D_8_, 98%) revealed a DCA-Phe conjugate with an [M+H]^+^ of *m/z* 540.3684 and a corresponding labeled ion with an [M+H]^+^ of *m/z* 548.4186, respectively (Figure 2C). Analysis of tandem MS fragmentation indicated condensation of the amine of Phe with DCA at the terminal side-chain carboxylic acid (Figure S3). Key MS^2^ fragments that indicate the position of the phenylalanine on DCA include the doubly dehydroxylated DCA-Phe conjugate ion (*m/z* 504.3472), the *N*-(1-Oxo-4-penten-1-yl)phenylalanine ion (*m/z* 248.1281), the phenylalanine aminium ion (*m/z* 166.0861), and the phenylalanine iminium ion (*m/z* 120.0805). Ions corresponding to the doubly dehydroxylated DCA-Phe conjugate ion and the phenylalanine aminium and iminium ions with eight Dalton increases were observed in MS^2^ data associated with an MS^1^ scan with an eight Dalton increase in the L-phenylalanine (D_8_, 98%) sample, further confirming the structural assignment of phenylalanine conjugation at the terminal side-chain carboxylic acid of DCA. Additionally, comparison of data from DCA-Phe in our study to that of phenylalanocholic acid (Phe-cholic acid) in Quinn, et al. (2) supports these results. Specifically, we observed matches between the phenylalanine ammonium and iminium ions as well as expected fragment shifts between the two MS^2^ spectra arising from the absence of a hydroxyl group in DCA-Phe.

#### Tyramine conjugate

Comparison of broth from P207 cultures containing DCA and unlabeled tyramine or isotopically labeled tyramine:HCl (1,1,2,2-D_4_, 98%) revealed a DCA-tyramine conjugate with an [M+H]^+^ of *m/z* 512.3740 and a corresponding labeled ion with an [M+H]^+^ of *m/z* 516.3987, respectively (Figure 2D). Analysis of tandem MS fragmentation indicated condensation of the amine of tyramine with DCA at the terminal side-chain carboxylic acid (Figure S4). Key MS^2^ fragments in data from the unlabeled P207 cultures that indicate the position of the tyramine on DCA include the doubly dehydroxylated DCA-tyramine conjugate ion (*m/z* 476.3526), the dehydroxylated DCA-tyramine conjugate ion (*m/z* 494.3629), the *N*-(1-Oxo-4-penten-1-yl)tyramine ion (*m/z* 220.1330), the tyramine cation (*m/z* 138.0913), and the 2-(4-hydroxyphenyl)ethylium ion (*m/z* 121.0644). Ions corresponding to these with four Dalton increases were observed in MS^2^ data associated with an MS^1^ scan with a four Dalton increase in the tyramine:HCl (1,1,2,2-D_4_, 98%) sample, further confirming the structural assignment of tyramine conjugation at the terminal side-chain carboxylic acid of DCA to produce tyraminodeoxycholic acid. The identity of the DCA-tyramine product was further validated by comparing the retention time and MS^2^ data of the species with an [M+H]^+^ of *m/z* 512.3740 to a DCA-tyramine synthetic standard. The matching retention times and MS_2_ data confirmed the identity of this molecule as DCA-tyramine (Figure 3 and Figure S5).

#### GABA conjugate

More specifically, comparison of broth from P207 cultures containing DCA and unlabeled 4-aminobutyric acid (GABA) or isotopically labeled GABA (^13^C_4_, 97-99%) revealed a DCA-GABA conjugate with an [M+H]^+^ of *m/z* 478.3531 and a corresponding labeled ion with an [M+H]^+^ of *m/z* 482.3666, respectively (Figure 2E). Analysis of tandem MS fragmentation indicated condensation of the primary amine of GABA with DCA at the terminal side-chain carboxylic acid (Figure S5). Key MS^2^ fragments in data from the unlabeled P207 cultures that indicate the position of GABA on DCA include the doubly dehydroxylated DCA-GABA conjugate ion (detected *m/z* 442.3316), the *N*-(1-Oxo-4-penten-1-yl)-GABA ion (detected *m/z* 186.1121), the GABA cation (detected *m/z* 104.0706), and the GABA iminium ion (detected *m/z* 86.0596). Samples from GABA (^13^C_4_, 97-99%)-labeled broth contained ions with four Dalton increases corresponding to each of these MS^2^ fragments, which were associated with an MS^1^ scan with a four Dalton increase. These results further support the structural assignment of GABA conjugation at the terminal side-chain carboxylic acid of DCA to produce γ-aminobutyrodeoxycholic acid.

**Table S1.** Postulated DCA-amine conjugates with predicted positive and negative ions used to search raw mass spectrometry data and molecular networks.

| **DCA-AA conjugate** | **[M+H]<sup>+</sup>** | **[M-H]<sup>-</sup>** |
| --- | --- | --- |
| DCA-Urea | 435.3218 | 433.3072 |
| DCA-Glycine | 450.3214 | 448.3068 |
| DCA-Glycolic acid | 451.3054 | 449.2908 |
| DCA-Alanine | 464.3371 | 462.3225 |
| DCA-beta-Alanine | 464.3371 | 462.3225 |
| DCA-Sarcosine | 464.3371 | 462.3225 |
| DCA-Lactic acid | 465.3211 | 463.3065 |
| DCA-Glycerol | 467.3367 | 465.3221 |
| DCA-Phosphoric acid | 473.2663 | 471.2517 |
| DCA-GABA | 478.3527 | 476.3381 |
| DCA-Serine | 480.3320 | 478.3174 |
| DCA-Uracil | 487.3167 | 485.3021 |
| DCA-ornithine lactam | 489.3687 | 487.3541 |
| DCA-Proline | 490.3527 | 488.3381 |
| DCA-Fumarate | 491.3004 | 489.2858 |
| DCA-Valine | 492.3684 | 490.3538 |
| DCA-Succinic acid | 493.3160 | 491.3014 |
| DCA-Threonine | 494.3476 | 492.3330 |
| DCA-Cysteine | 496.3092 | 494.2946 |
| DCA-Nicotinamide | 497.3374 | 495.3228 |
| DCA-Taurine | 500.3041 | 498.2895 |
| DCA-5-oxoprolinate | 504.3320 | 502.3174 |
| DCA-5-oxoproline | 504.3320 | 502.3174 |
| DCA-L-Isoleucine | 506.2894 | 504.2748 |
| DCA-Leucine | 506.3840 | 504.3694 |
| DCA-Isoleucine | 506.3840 | 504.3694 |
| DCA-Asparagine | 507.3429 | 505.3283 |
| DCA-ornithine | 507.3793 | 505.3647 |
| DCA-Aspartic acid | 508.3269 | 506.3123 |
| DCA-Malate | 509.3109 | 507.2963 |
| DCA-Erythronic acid | 511.3266 | 509.3120 |
| DCA-Hypoxanthine | 511.3279 | 509.3133 |
| DCA-tyramine | 512.3735 | 510.3589 |
| DCA-2-Oxoglutarate | 521.3109 | 519.2963 |
| DCA-Glutamine | 521.3585 | 519.3439 |
| DCA-Lysine | 521.3949 | 519.3803 |
| DCA-Glutamic acid | 522.3426 | 520.3280 |
| DCA-Methionine | 524.3405 | 522.3259 |
| DCA-histidine | 530.3589 | 528.3443 |
| DCA-Phenylalanine | 540.3684 | 538.3538 |
| DCA-selenocysteine | 543.2458 | 541.2312 |
| DCA-Capric acid | 547.4357 | 545.4211 |
| DCA-arginine | 549.4011 | 547.3865 |
| DCA-N-Acetylaspartic acid | 550.3375 | 548.3229 |
| DCA-Tyrosine | 556.3633 | 554.3487 |
| DCA-Azelaic acid | 563.3943 | 561.3797 |
| DCA-N-Acetylglutamic acid | 564.3531 | 562.3385 |
| DCA-Citrate | 567.3164 | 565.3018 |
| DCA-Isocitrate | 567.3164 | 565.3018 |
| DCA-tryptophan | 579.3793 | 577.3647 |
| DCA-Pantothenic acid | 594.3994 | 592.3848 |
| DCA-Myristic acid | 603.4983 | 601.4837 |
| DCA-Palmitoleic acid | 629.5140 | 627.4994 |
| DCA-pyrollysine | 630.4477 | 628.4331 |
| DCA-Palmitic acid | 631.5296 | 629.5150 |
| DCA-Oleic acid | 657.5453 | 655.5307 |
| DCA-Stearic acid | 659.5609 | 657.5463 |
| DCA-Docosahexaenoic acid | 703.5296 | 701.5150 |
| DCA-Cholesterol | 761.6443 | 759.6297 |
| DCA-(2R)-Pyrrolidine | 762.9894 | 760.9748 |

**Figure S1.**
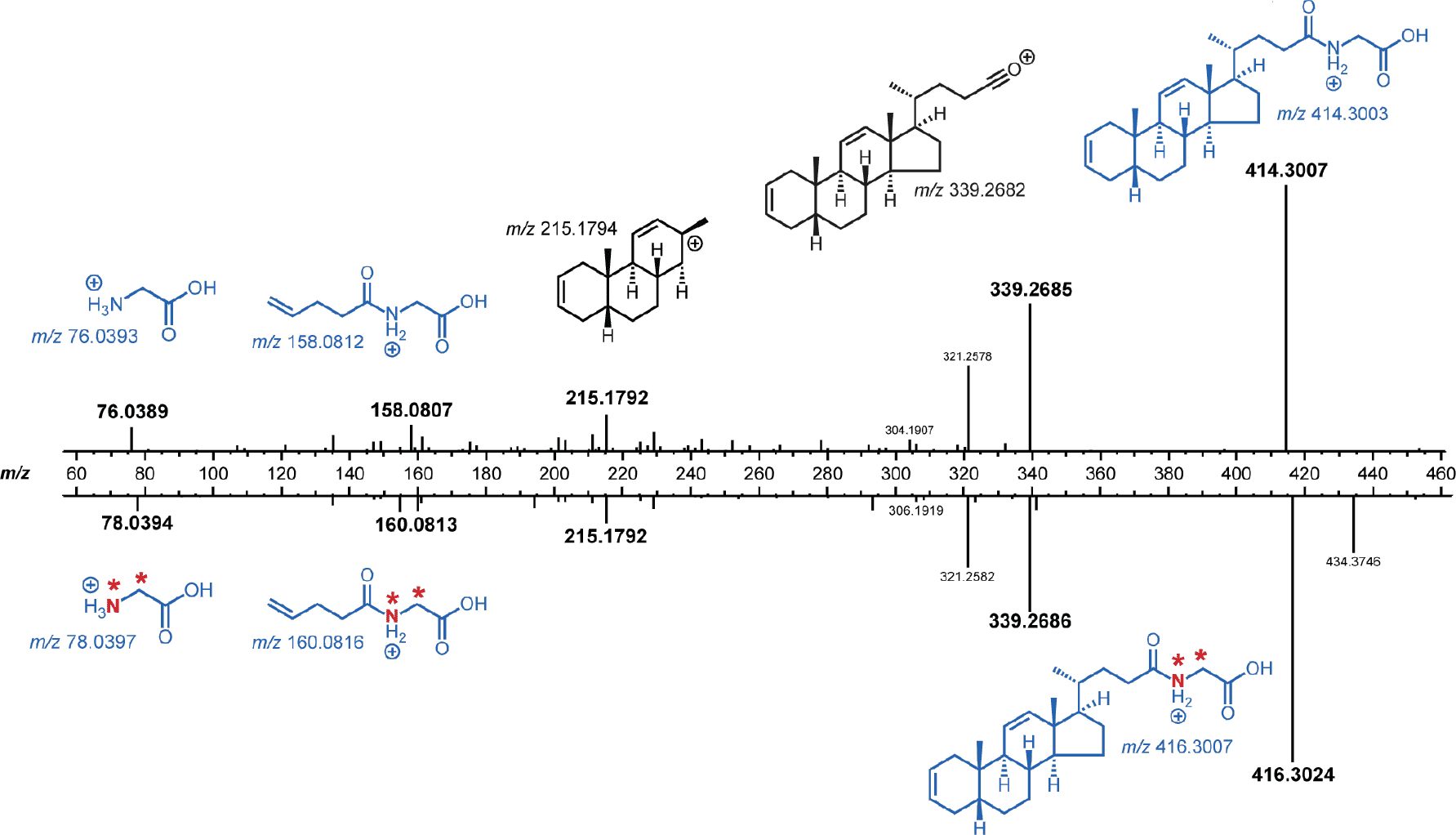
Mirror plot of a summed DCA-Gly (*m/z* 450.3220) MS^2^ spectrum opposite a summed MS^2^ spectrum for the isotopically labeled DCA-^15^N,^13^C-Gly (*m/z* 452.3230), which was observed only when *B. fragilis* P207 was cultured in the presence of DCA and ^15^N,^13^C-Gly (Figure 2). The +2 deuterium shift was observed in the expected MS^2^ fragments between the spectra. (red asterisk indicating location of deuteriums). All structural assignments for MS2 ions are putative.

**Figure S2.**
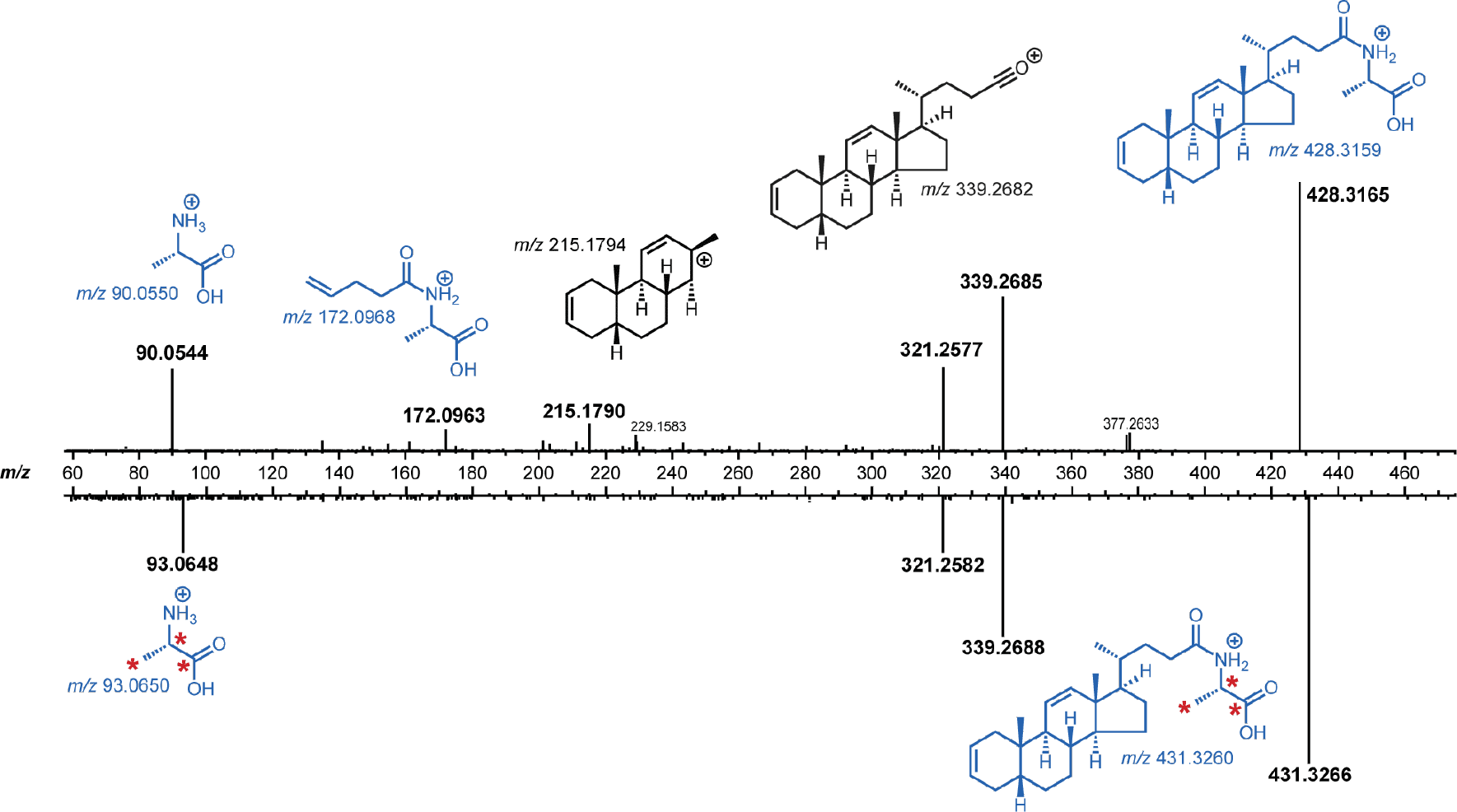
Mirror plot of a summed DCA-Ala (*m/z* 464.3373) MS^2^ spectrum opposite a summed MS^2^ spectrum for the isotopically labeled DCA-^13^C_3_-Ala (*m/z* 467.3475), which was observed only when *B. fragilis* P207 was cultured in the presence of DCA and ^13^C_3_-Ala (Figure 2). The +3 deuterium shift was observed in the expected MS^2^ fragments between the spectra (red asterisk indicating location of deuteriums). All structural assignments for MS2 ions are putative.

**Figure S3.**
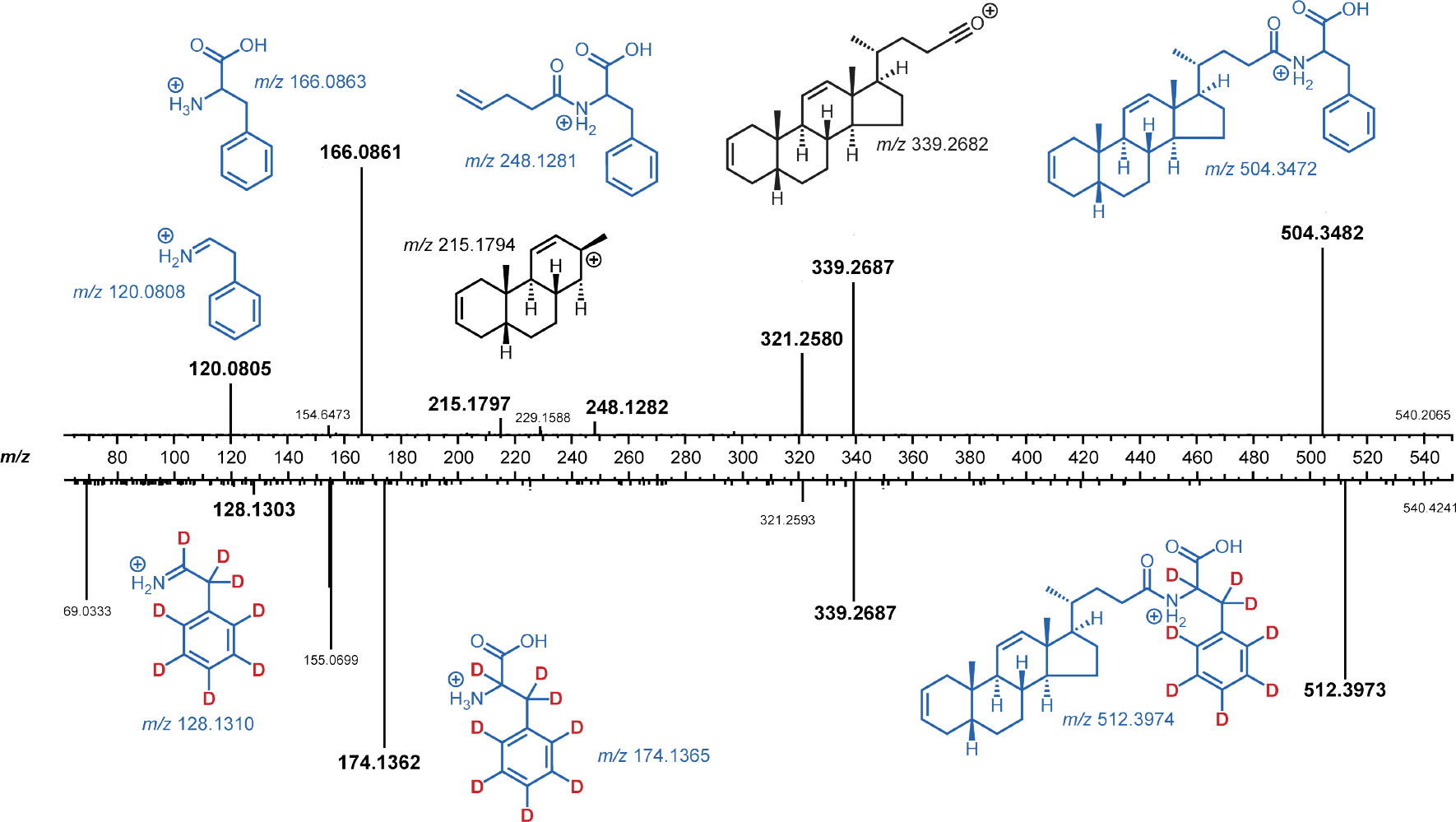
Mirror plot of a summed DCA-Phe (*m/z* 540.3686) MS^2^ spectrum opposite a summed MS^2^ spectrum for the isotopically labeled DCA-D_8_-Phe (*m/z* 548.4186), which was observed only when *B. fragilis* P207 was cultured in the presence of DCA and D_8_-Phe (Figure 2). The +8 deuterium shift was observed in the expected MS^2^ fragments between the spectra (red text indicating location of deuteriums). All structural assignments for MS2 ions are putative.

**Figure S4.**
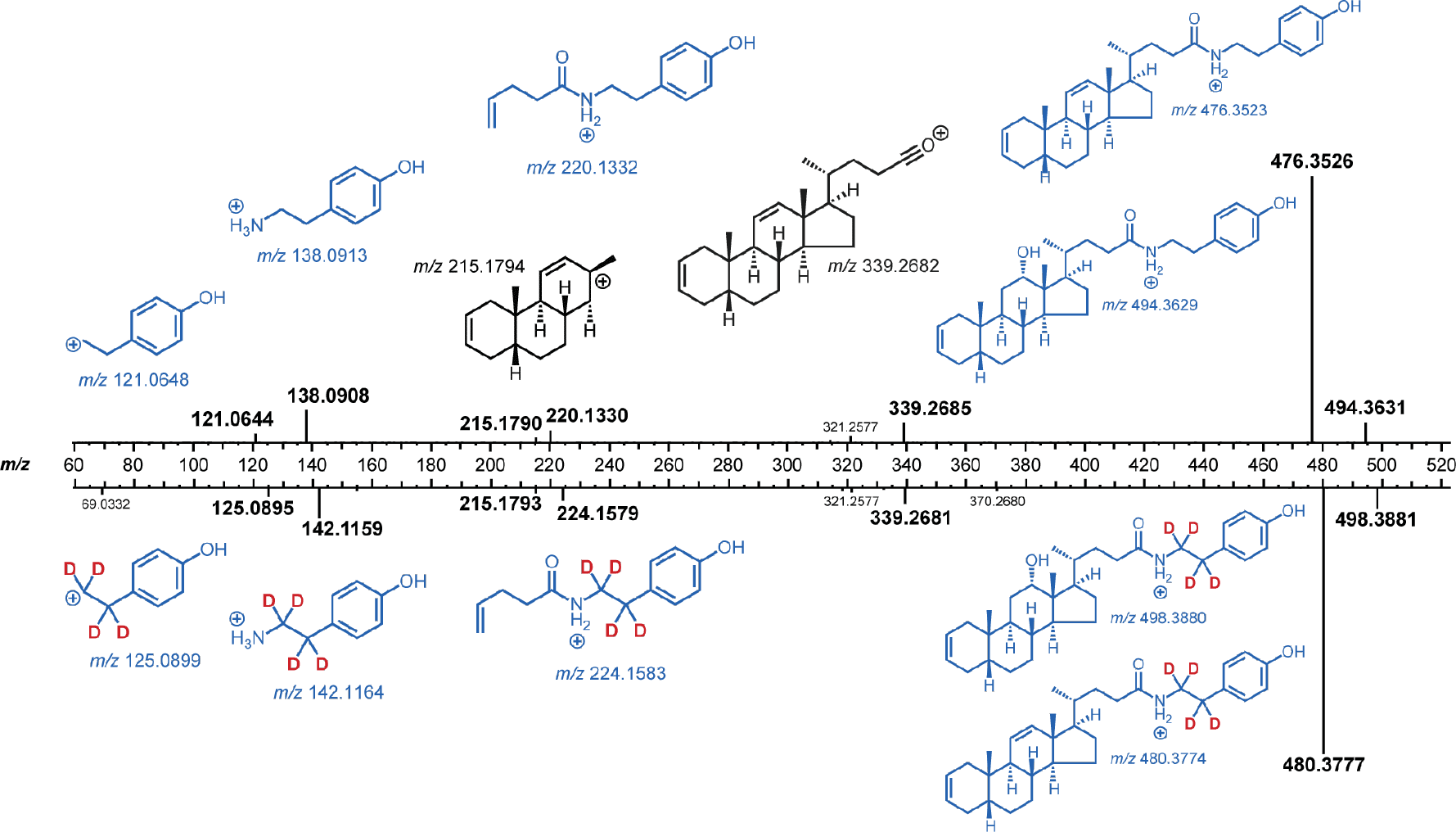
Mirror plot of a summed DCA-tyramine (*m/z* 512.3740) MS^2^ spectrum opposite a summed MS^2^ spectrum for the isotopically labeled DCA-D_4_-tyramine (*m/z* 516.3987), which was observed only when *B. fragilis* P207 was cultured in the presence of DCA and D_4_-tyramine (Figure 2). The +4 deuterium shift was observed in the expected MS^2^ fragments between the spectra (red asterisk indicating location of deuteriums). All structural assignments for MS2 ions are putative.

**Figure S5.**
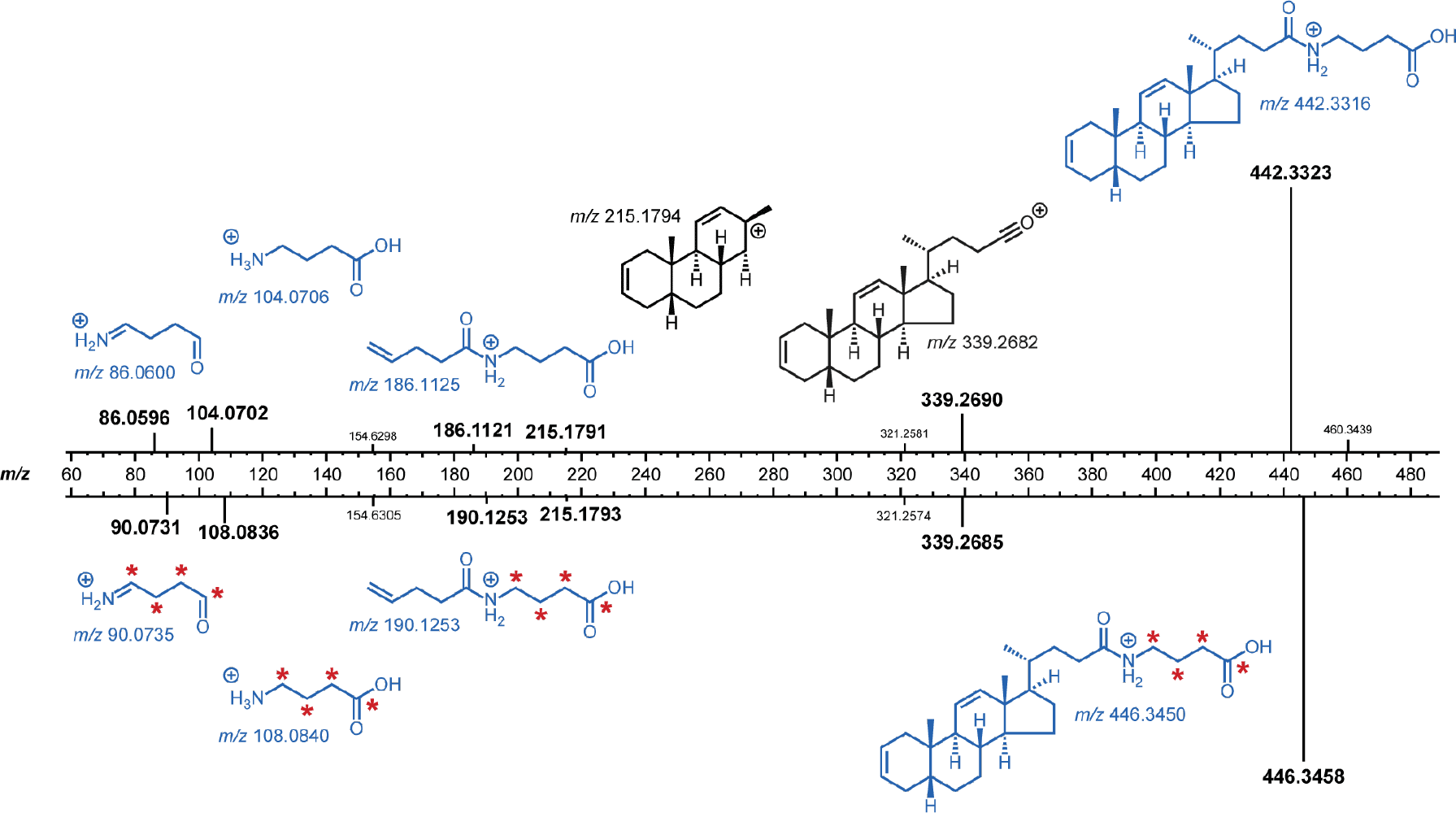
Mirror plot of a summed DCA-GABA (*m/z* 478.3531) MS^2^ spectrum opposite a summed MS^2^ spectrum for the isotopically labeled DCA-^13^C_4_-GABA (*m/z* 482.3666), which was observed only when *B. fragilis* P207 was cultured in the presence of DCA and ^13^C_4_-GABA (Figure 2). The +4 deuterium shift was observed in the expected MS^2^ fragments between the spectra (red asterisk indicating location of deuteriums). All structural assignments for MS2 ions are putative.

**Figure S6.**
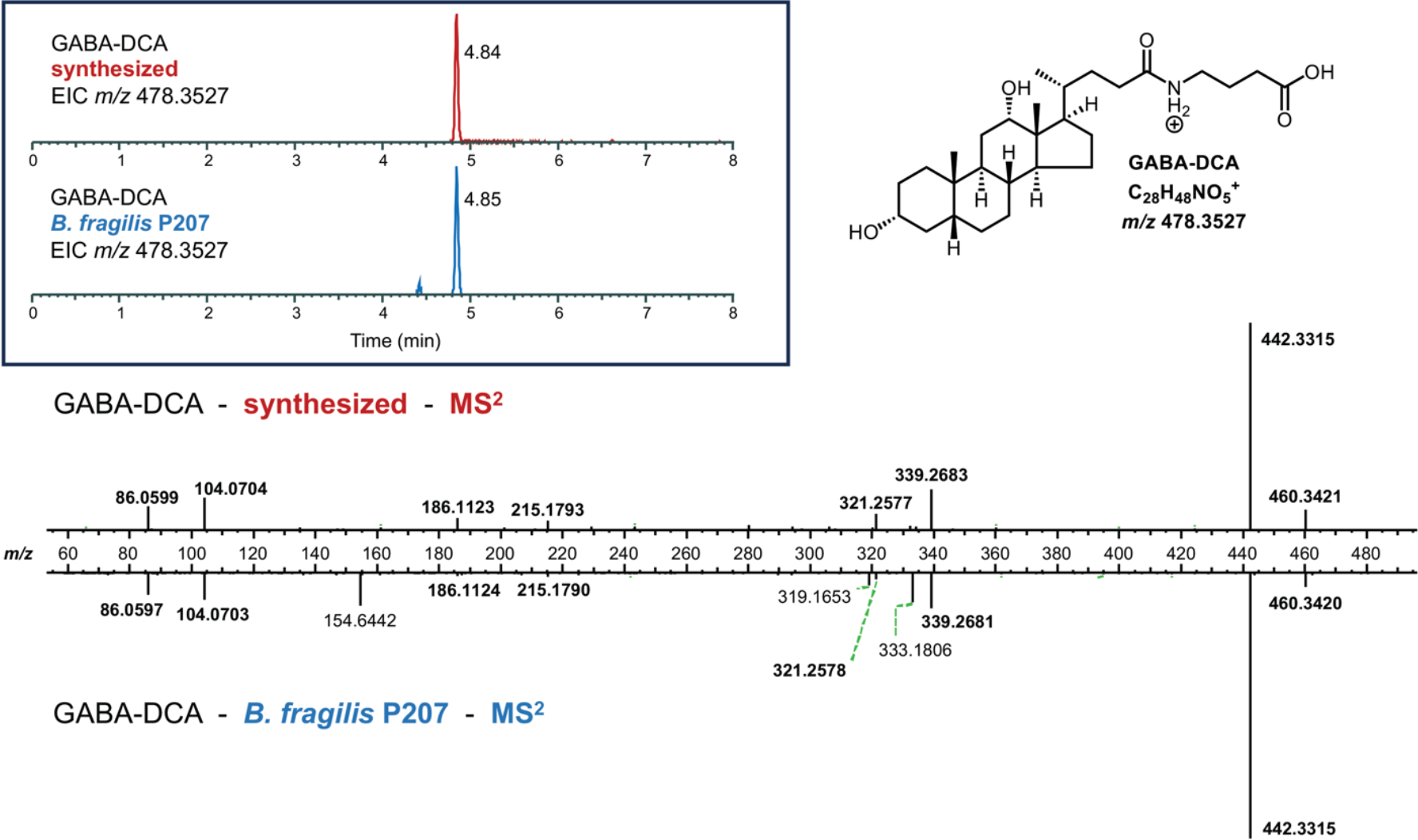
Comparative LC-MS/MS analysis of DCA-GABA produced by *B. fragilis* P207 *in vitro* to a synthetic standard. (Top left) Extracted ion chromatogram (EIC) of chemically synthesized DCA-GABA (m/z 478.3527) (retention time=4.84 minutes) and the EIC for DCA-GABA identified in *B. fragilis* P207 broth cultures (bottom; retention time=4.85 minutes). (bottom) Mirror plot of a summed DCA-GABA MS^2^ spectrum for the synthetic standard opposite a summed MS^2^ spectrum for the DCA-GABA detected in *B. fragilis* P207 broth cultures.

**Figure S7.**
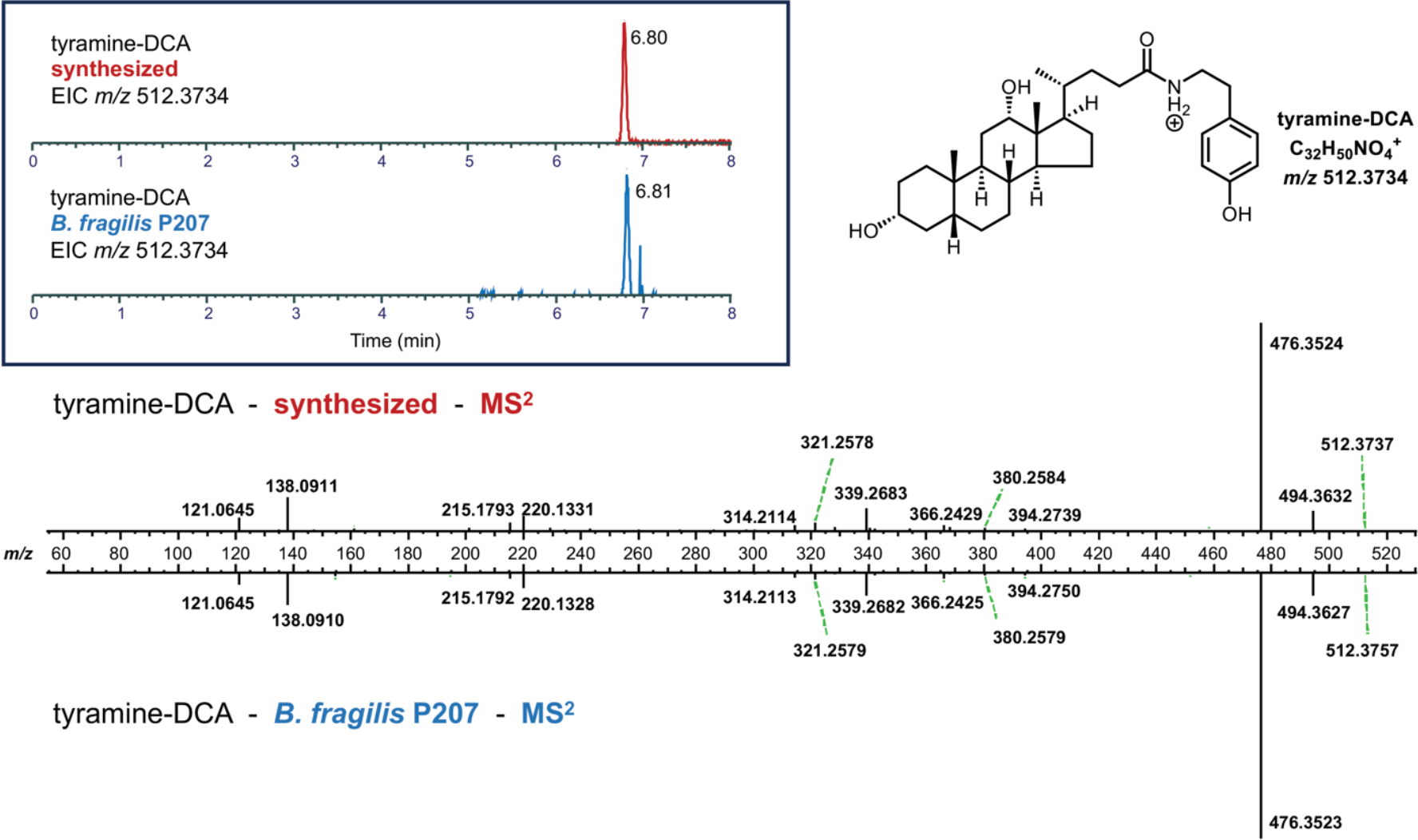
Comparative LC-MS/MS analysis of DCA-tyramine produced by *B. fragilis* P207 *in vitro* to a synthetic standard. (Top left) Extracted ion chromatogram (EIC) of chemically synthesized DCA-tyramine (m/z 512.3734) (retention time=6.80 minutes) and the EIC for DCA-tyramine identified in *B. fragilis* P207 broth cultures (bottom; retention time=6.81 minutes). (bottom) Mirror plot of a summed DCA-GABA MS^2^ spectrum for the synthetic standard opposite a summed MS^2^ spectrum for the DCA-GABA detected in *B. fragilis* P207 broth cultures.

**Figure S8.**
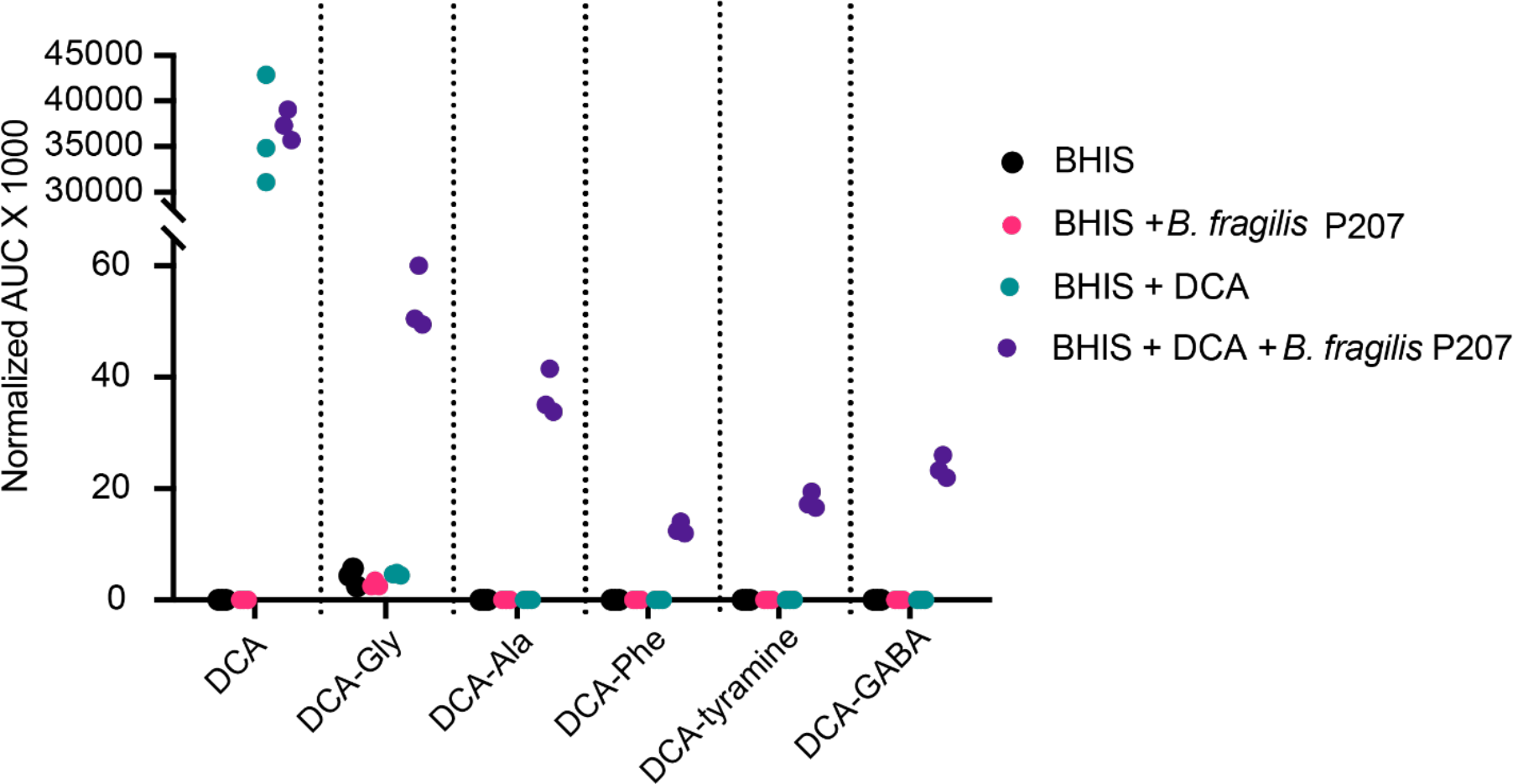
Relative abundances of the five deoxycholic acid (DCA)-amine conjugates in triplicate samples of BHIS media, BHIS media with *B. fragilis* strain P207 (P207), BHIS media with DCA, and BHIS media with P207 and DCA normalized to the average peak area under the curve (AUC) for internal standard ions D_4_-taurocholic acid [M+H]^+^ at *m/z* 520.3241, D_4_-taurodeoxycholic acid [M+H]^+^ at *m/z* 504.3291, and D_4_-glycocholic acid at m/z 470.3414.

**Figure S9.**
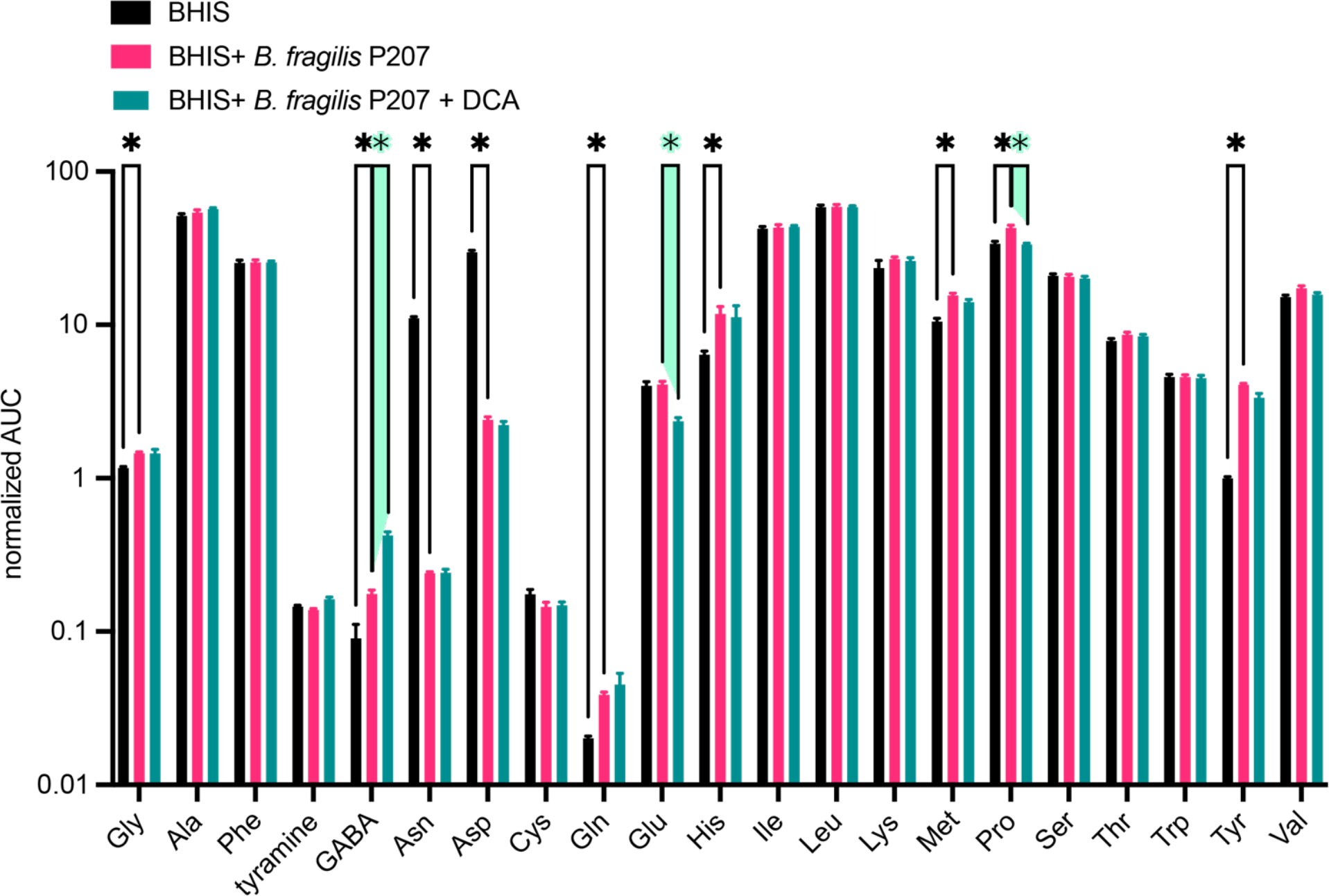
Comparative GC/MS-based quantification of select amines in *B. fragilis* P207 cultures. The cultures were incubated in BHIS medium either without (blue bars) or with (pink bars) the addition of deoxycholic acid (DCA) at a concentration of 0.01% (w/v). BHIS medium alone is shown in black bars. The graph presents the relative concentrations, derived from the area under the curve (AUC) of detected peaks, indicating the comparative levels of these amines across the different conditions. Each bar denotes the mean of three independent experiments, and error bars represent the standard deviations. Statistical significance between conditions was assessed using individual t-tests, with corrections for multiple comparisons via the two-stage step-up method of Benjamini, Krieger, and Yekutieli; significance is indicated by an asterisk (*), denoting a q-value of less than 0.01.

**Figure S10.**
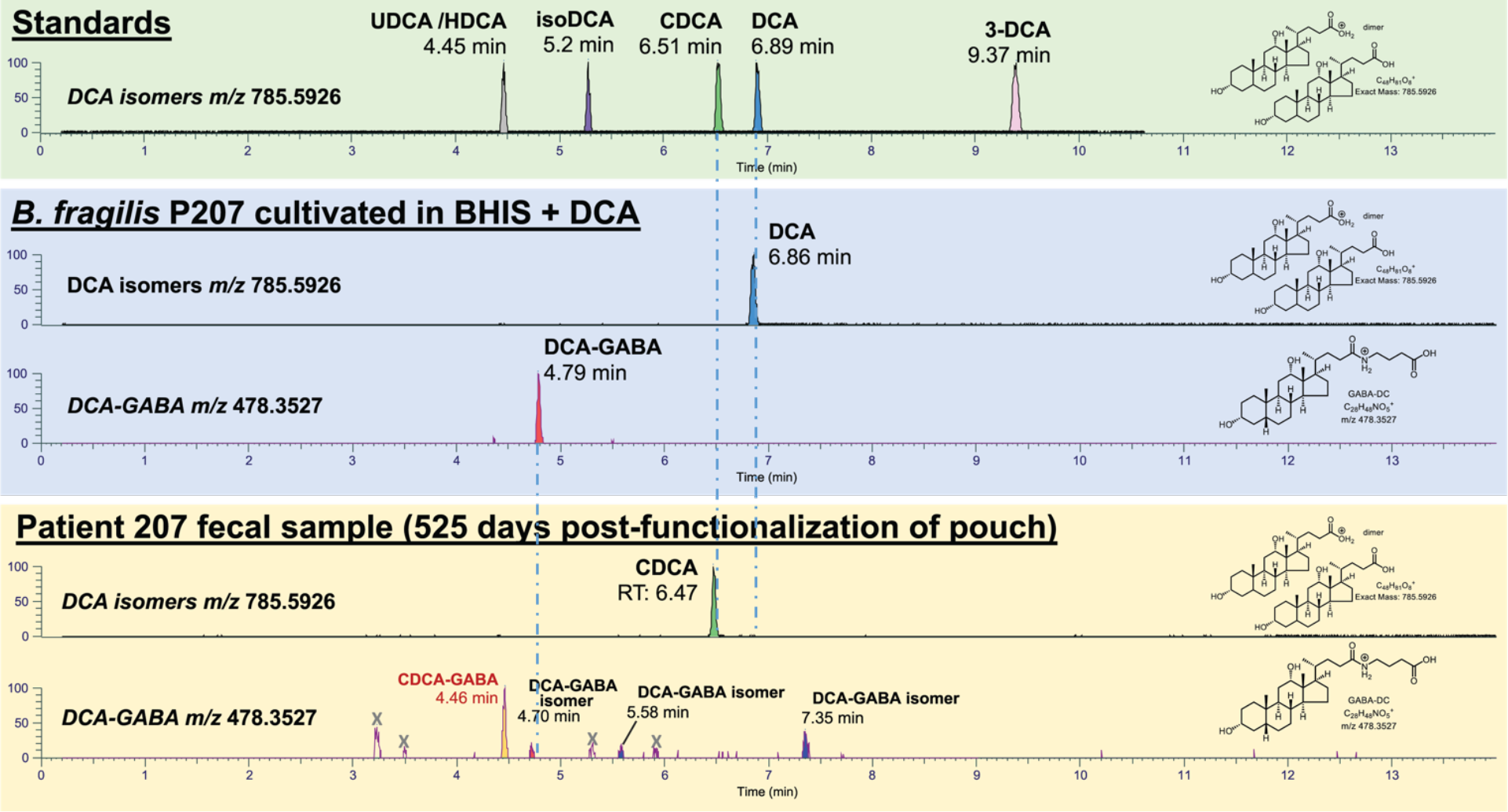
(Top panel) Liquid chromatography temporal elution profile of deoxycholic acid (DCA) and DCA isomer standards (*m/z* dimer 785.5926): ursodeoxycholic acid (UDCA); hyodeoxycholic acid (HDCA); chenodeoxycholic acid (CDCA). (Middle panel) Extracted ion chromatogram of resolved BHIS media containing *B. fragilis* strain P207 with 0.01% (w/v) DCA; *m/z* for DCA (dimer) and DCA-GABA. Quantification of all five DCA conjugates detected in *B. fragilis* strain P207 culture extract by MS/MS are presented in Figure S8 and Table S2. (bottom panel) Extracted ion chromatogram of fecal extract of human pouchitis patient 207 at 525 days post pouch functionalization; *m/z* for DCA (dimer) and DCA-GABA. This analysis was conducted on multiple fecal collection time points and yielded multiple peaks with MS/MS profiles matching GABA conjugated to DCA. The absence of a clear peak at Rt ≈4.8 minutes (corresponding to the DCA-GABA standard; see Figure 3), and the lack of DCA in patient 207 stool (Rt ≈ 6.9) suggests these species are DCA isomer-GABA conjugates (labeled DCA-GABA isomer). LC-MS/MS analysis of a synthetic GABA-CDCA standard showed that the major GABA conjugate peak at this *m/z* is CDCA-GABA. Quantification of bile acid peaks in longitudinal fecal samples across an approximate 2-year period after surgical functionalization of the ileal pouch is presented in Table S2.

**Figure S11.**
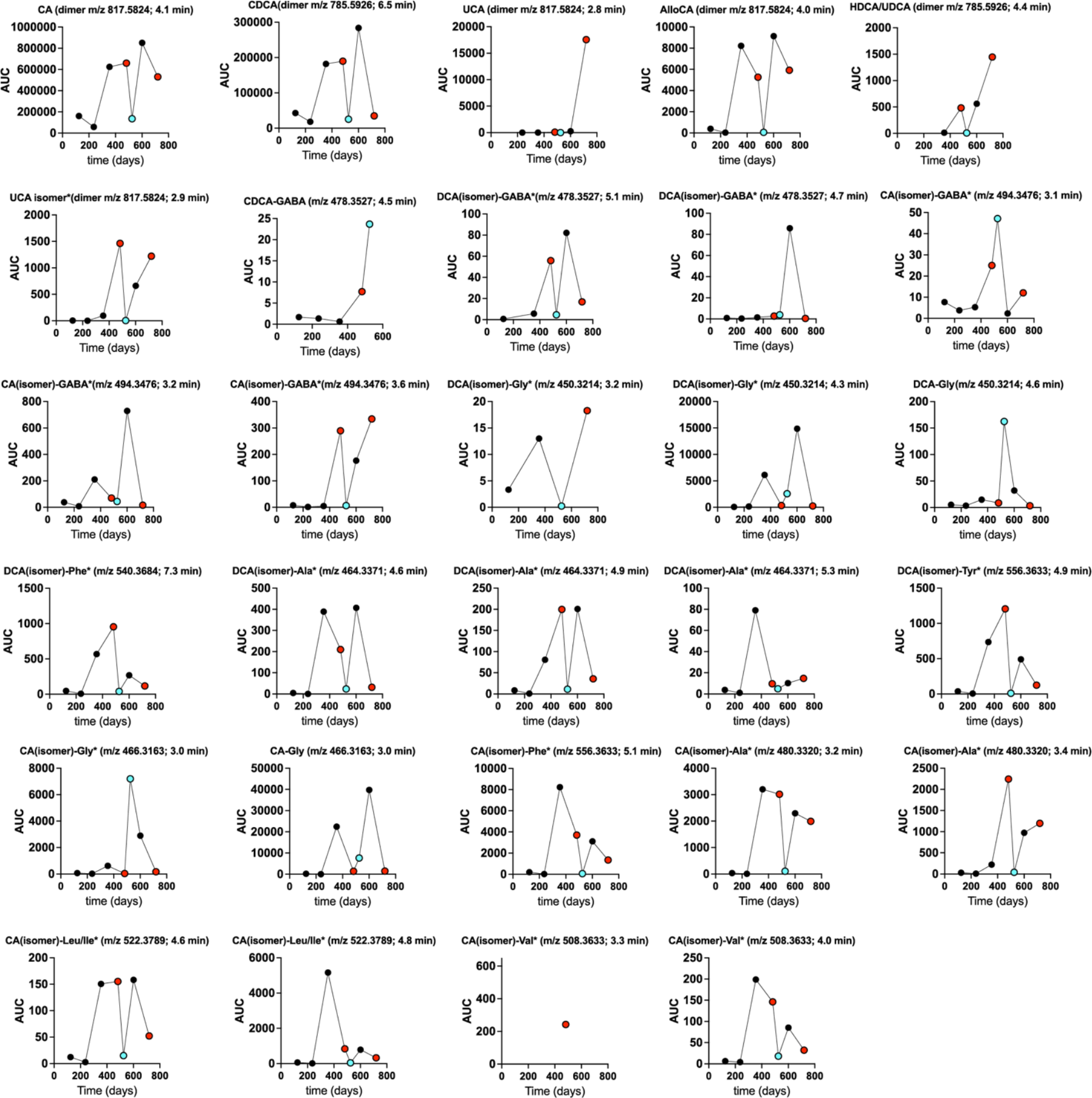
Longitudinal profile of unconjugated and conjugated bile acid species in pouchitis patient 207 from 124 to 719 days post-functionalization of the ileal J-pouch (3). Red dots at 482 and 719 days post-functionalization are clinic visits where the patient was diagnosed with pouch inflammation (i.e. pouchitis). Blue dot (525 days) was during a period of antimicrobial treatment (ciprofloxacin), in which the patient did not have pouchitis. The top of each graph notes the elution time of each species in minutes. Synthetic standards of hyodeoxycholate/ursodeoxycholate (HDCA/UDCA), chenodeoxycholate (CDCA), cholic acid (CA), deoxycholic acid (DCA), 3-CDA, iso-DCA, ursocholic acid (UCA), and litchcholic acid (LCA) were included in this analysis. DCA (3α,12α-Dihydroxy-5β-cholan-24-oic acid) was absent at all time points in this patient, so we expect the labeled DCA(isomer) amide conjugates have a CDCA core, though this is not proven in most cases. Likewise, we expect that many of the abundant CA(isomer) conjugates have a cholic acid core though the particular isomer/epimer of these products has not been defined. Points are missing for compounds at timepoints where they were not detected.

**Figure S12.**
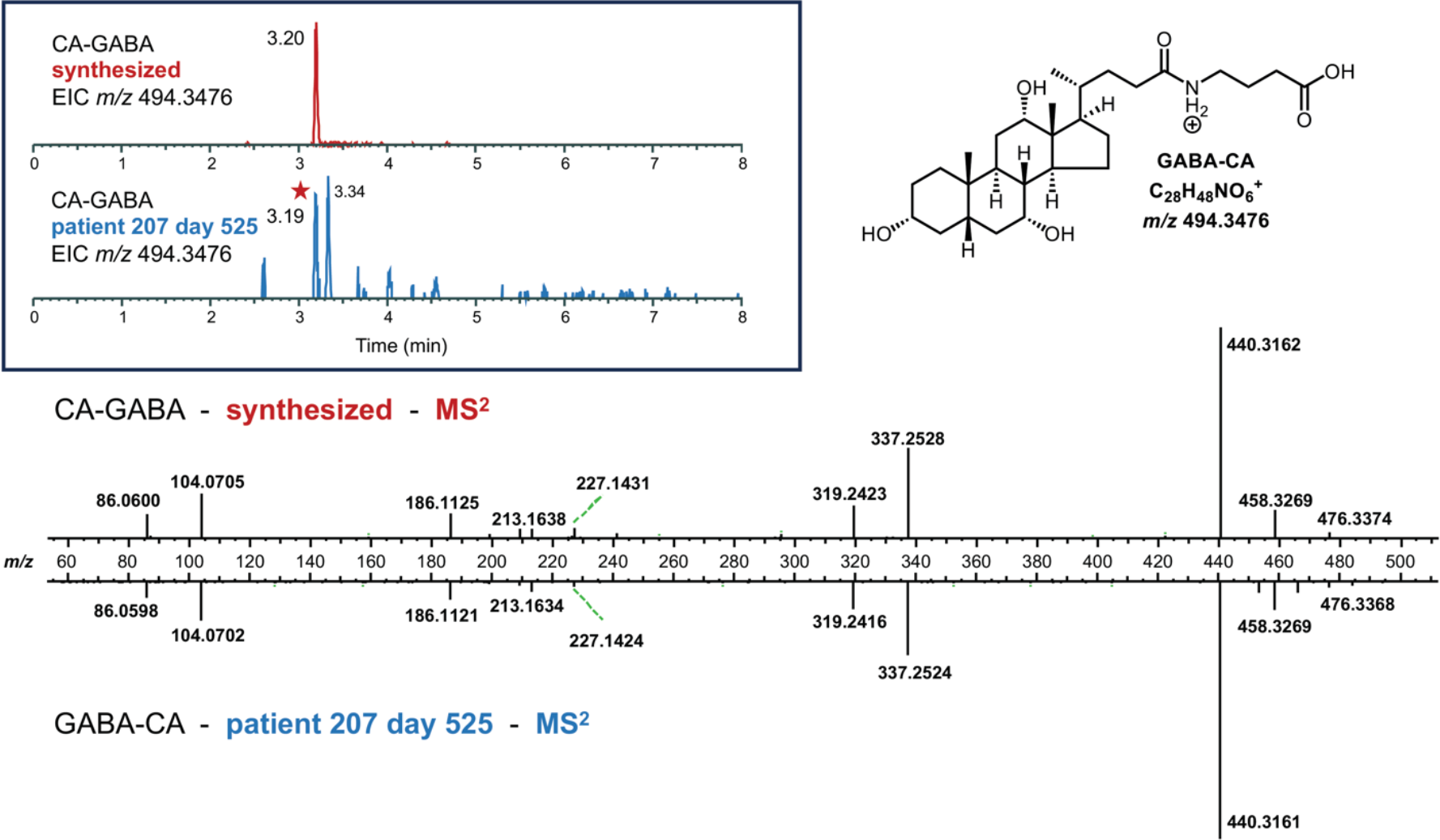
Comparative LC-MS/MS analysis of CA-GABA in patient 207 stool to a synthetic standard. (Top left) Extracted ion chromatogram (EIC) of chemically synthesized CA-GABA (*m/z* 494.3476) (retention time=3.20 minutes) and the EIC for CA-GABA identified in patient 207 stool at 525 days post-functionalization of the ileal pouch (bottom; retention time=3.19 minutes). (bottom) Mirror plot of a summed CA-GABA MS^2^ spectrum for the synthetic standard opposite a summed MS^2^ spectrum for the CA-GABA detected in patient 207 stool.

**Figure S13.**
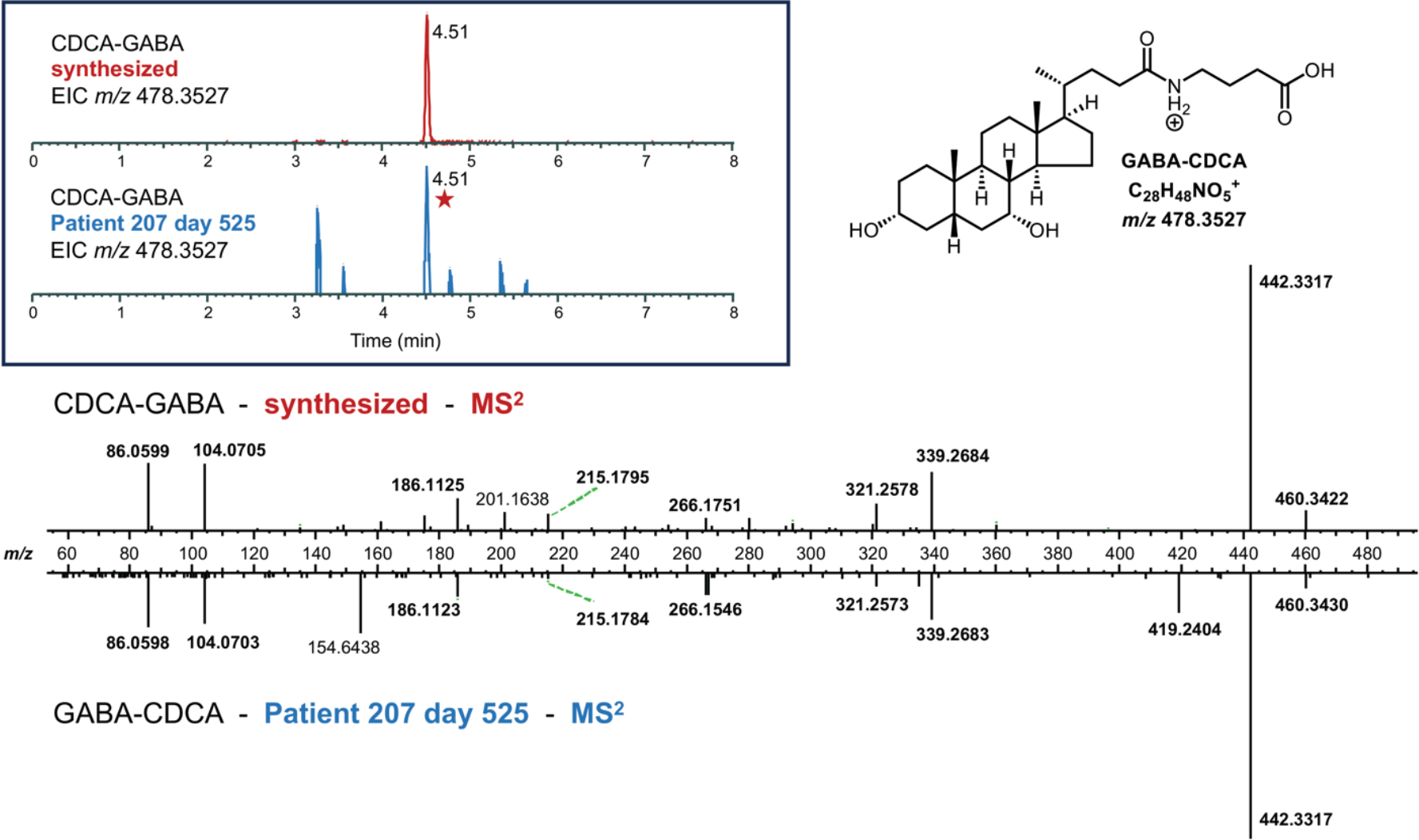
Comparative LC-MS/MS analysis of CDCA-GABA in patient 207 stool to a synthetic standard. (Top left) Extracted ion chromatogram (EIC) of chemically synthesized CDCA-GABA (*m/z* 478.3527) (retention time=4.51 minutes) and the EIC for CDCA-GABA identified in patient 207 stool at 525 days post-functionalization of the ileal pouch (bottom; retention time=4.51 minutes). (bottom) Mirror plot of a summed CDCA-GABA MS^2^ spectrum for the synthetic standard opposite a summed MS^2^ spectrum for the CDCA-GABA detected in patient 207 stool.

**Figure S14.**
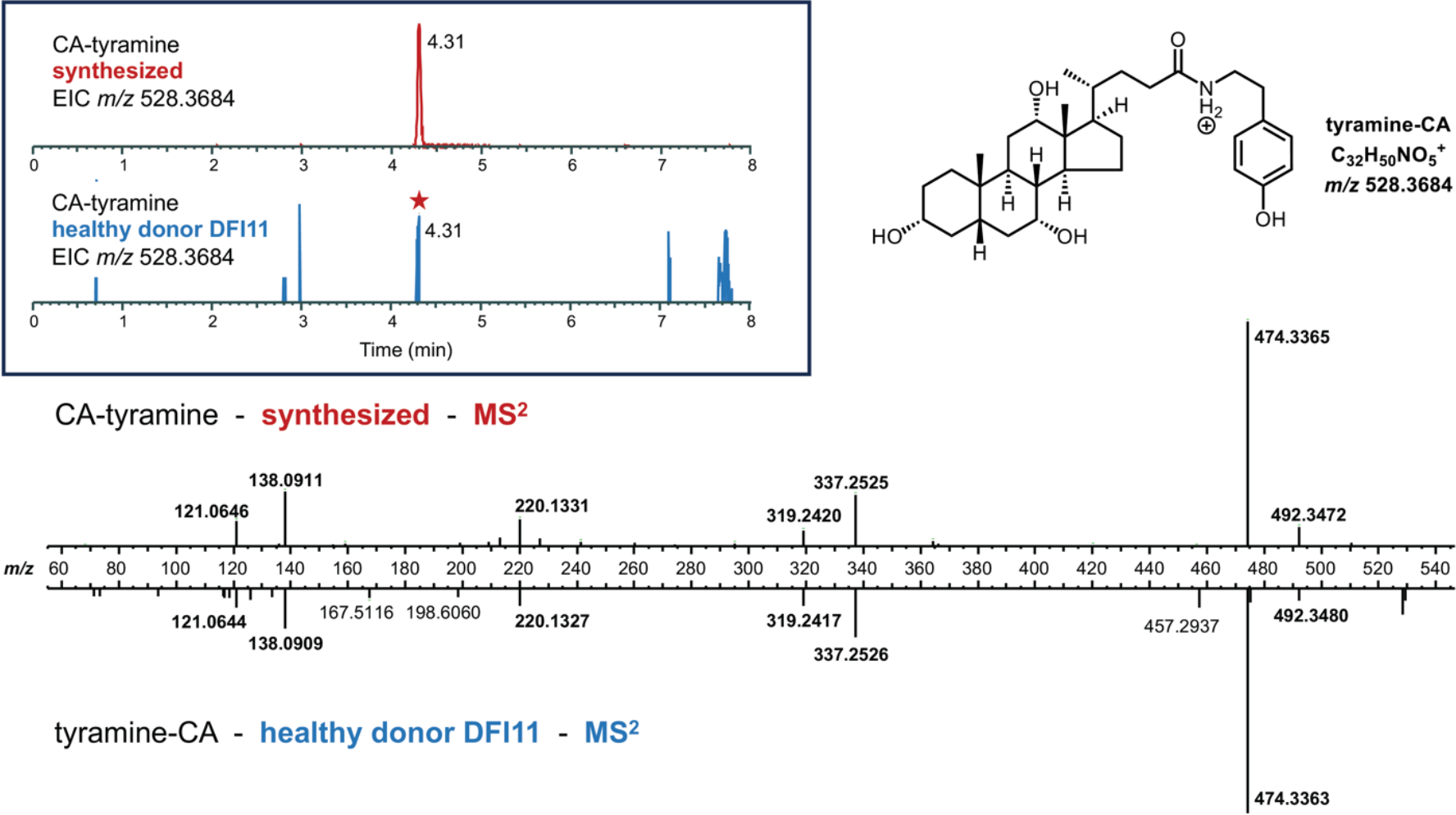
Comparative LC-MS/MS analysis of CA-tyramine in healthy human donor (DFI11) stool to a synthetic standard. (Top left) Extracted ion chromatogram (EIC) of chemically synthesized CA-tyramine (*m/z* 528.3684) (retention time=4.31 minutes) and the EIC for CA-tyramine identified in healthy donor DFI11 stool (bottom; retention time=4.31 minutes). (bottom) Mirror plot of a summed CA-tyramine MS^2^ spectrum for the synthetic standard opposite a summed MS^2^ spectrum for the CA-tyramine in DFI11 donor stool.

**Figure S15.**
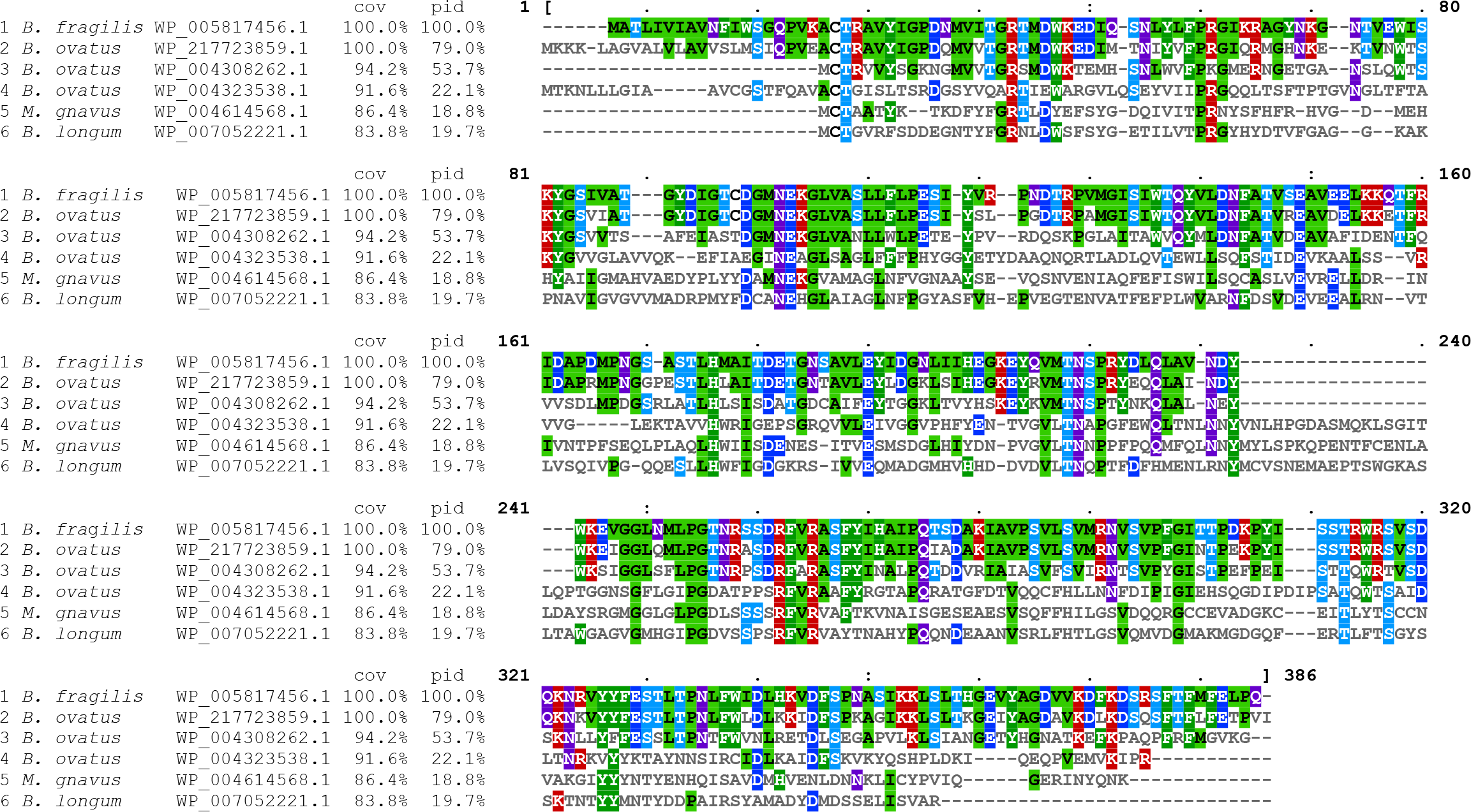
Multiple sequence alignment of N-terminal nucleophilic cysteine hydrolase (Ntn) enzymes (Conserved Domain Database accession cd01902) from *B. fragilis* P207 (reference sequence for this alignment; WP_005817456.1), *Mediterraneibacter gnavus* MSK15.77 (WP_004614568.1), *Bifidobacterium longum* DFI.2.45 (WP_007052221.1), and *Bacteroides ovatus* MSK22.29 (which encodes three Ntn paralogs: WP_217723859.1, WP_004308262.1, and WP_004323538.1). cov-percent coverage; pid-percent identity.

**Figure S16.**
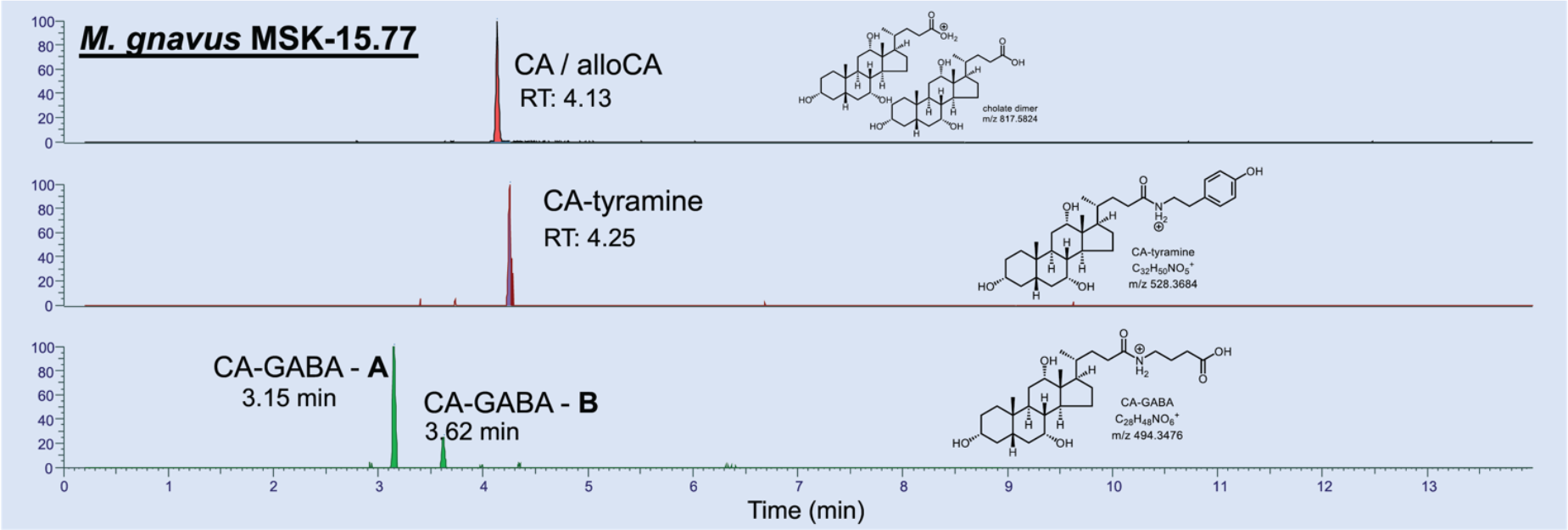
Incubation of *M. gnavus* strain MSK-15.77 with 0.01% (w/v) cholic acid (CA) in BHIS broth followed by separation of culture extract by liquid chromatography provides evidence for production of CA-tyramine and two species with MS/MS profiles matching CA-GABA (putative isomers).

**Figure S17.**
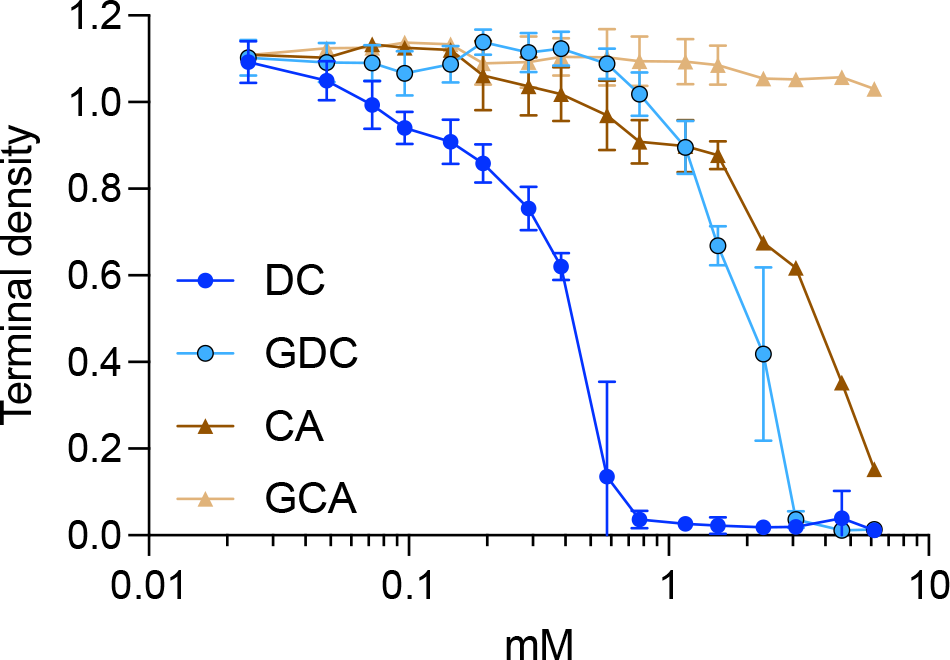
Glycine conjugation to DCA and CA reduces toxicity to *B. fragilis* P207 *in vitro*. Terminal density measurements (OD_600_) of *B. fragilis* P207 (24 hours of growth) in BHIS under increasing concentrations of bile acids (DC-deoxycholic acid; GDC-glycodeoxycholic acid; CA-cholic acid; GCA-glycocholic acid). Data show five biological replicates for DC and CA, and three replicates for GDC and GCA; error bars represent standard deviation.

